# Dysregulation of external globus pallidus-subthalamic nucleus network dynamics in Parkinsonian mice

**DOI:** 10.1101/774091

**Authors:** Ryan F. Kovaleski, Joshua W. Callahan, Marine Chazalon, Jérôme Baufreton, Mark D. Bevan

## Abstract

Reciprocally connected GABAergic external globus pallidus (GPe) and glutamatergic subthalamic nucleus (STN) neurons form a key, centrally-positioned network within the basal ganglia, a group of subcortical brain nuclei critical for voluntary movement. In Parkinson’s disease (PD) and its models, abnormal rates and patterns of GPe-STN network activity are linked to motor dysfunction. Using cell class-specific optogenetic identification and inhibition approaches during cortical slow-wave activity and activation, we report that in dopamine-depleted mice 1) D2 dopamine receptor expressing striatal projection neurons (D2-SPNs) are hyperactive 2) prototypic parvalbumin (PV)-expressing GPe neurons are excessively patterned by D2-SPNs 3) despite being disinhibited, STN neurons are not hyperactive 4) the STN opposes rather than facilitates abnormal striatopallidal patterning. Together with recent studies, these data argue that in Parkinsonian mice abnormal, temporally offset PV GPe neuron and STN activity results from increased striatopallidal transmission and that compensatory plasticity within the STN prevents its hyperactivity.

## Introduction

Reciprocally connected GABAergic external globus pallidus (GPe) and glutamatergic subthalamic nucleus (STN) neurons form a key, centrally-located network within the basal ganglia, a group of subcortical brain nuclei critical for the execution of motivated, goal-directed, and habitual behaviors (Keeler, Pretsell, & Robbins, 2014; Klaus, Alves da Silva, & Costa, 2019; Mink & Thach, 1993; Nelson & Kreitzer, 2014; Redgrave et al., 2010; Yin & Knowlton, 2006). The GPe and STN are components of the so-called “indirect pathway”, which emanates from D2 receptor expressing GABAergic striatal projection neurons (D2-SPNs), traverses the GPe and STN, and terminates in the GABAergic basal ganglia output nuclei, the internal globus pallidus (GPi), and substantia nigra pars reticulata (SNr) (Albin, Young, & Penney, 1989; Gerfen et al., 1990). Cortical (or thalamic) activation of D2-SPNs leads through the indirect pathway to increased inhibition of basal ganglia targets, which was traditionally thought to suppress movement (Albin et al., 1989; Frank, Seeberger, & O’Reilly R, 2004; Kravitz et al., 2010; Maurice, Deniau, Glowinski, & Thierry, 1999; Tachibana, Kita, Chiken, Takada, & Nambu, 2008). Indeed, sustained optogenetic activation of D2-SPNs rapidly arrests motion (Kravitz et al., 2010). However, recent studies argue that indirect pathway activity is necessary for the efficient execution of action sequences, possibly through the suppression of competing actions and termination of selected actions. Thus, D2-SPNs are recruited during natural and trained behaviors, and optogenetically inhibiting them interrupts execution of action sequences (Barbera et al., 2016; Carvalho Poyraz et al., 2016; Cui et al., 2013; Lambot et al., 2016; LeBlanc et al., 2018; Lemos et al., 2016; Markowitz et al., 2018; Sippy, Lapray, Crochet, & Petersen, 2015; Tecuapetla, Jin, Lima, & Costa, 2016). The STN is also a component of the so-called hyperdirect pathway, which rapidly relays cortical excitation to the basal ganglia output nuclei (and GPe) (Degos, Deniau, Le Cam, Mailly, & Maurice, 2008; T. Kita & Kita, 2012; Maurice et al., 1999; Nambu, Takada, Inase, & Tokuno, 1996; Nambu, Tokuno, & Takada, 2002; Tachibana et al., 2008). This pathway is also critical for motor control, possibly through the rapid constraint of competing actions and/or stalling of movement during decision making (Aron & Poldrack, 2006; Baunez, Nieoullon, & Amalric, 1995; Engel & Fries, 2010; Isoda & Hikosaka, 2008; Jahanshahi, Obeso, Rothwell, & Obeso, 2015; Mink & Thach, 1993; Nambu et al., 2002; Schmidt, Leventhal, Mallet, Chen, & Berke, 2013; Zavala, Zaghloul, & Brown, 2015).

In Parkinson’s disease (PD), abnormal GPe-STN network activity is linked to the expression of motor symptoms, specifically akinesia, bradykinesia, and rigidity (Albin et al., 1989; Hammond, Bergman, & Brown, 2007; McGregor & Nelson, 2019; Quiroga-Varela, Walters, Brazhnik, Marin, & Obeso, 2013). The classic view is that loss of dopamine leads to elevation of D2-SPN activity, which leads to increased inhibition of GPe neurons, disinhibition of STN neurons, and increased basal ganglia output, which suppresses movement (Albin et al., 1989; Kravitz et al., 2010; Mink & Thach, 1993). However, firing rate changes in the Parkinsonian basal ganglia are often moderate in nature or absent, and in some cases opposite to those predicted by the classic model (Bergman, Wichmann, Karmon, & DeLong, 1994; Goldberg et al., 2002; Leblois, Boraud, Meissner, Bergman, & Hansel, 2006; McGregor & Nelson, 2019; Nelson & Kreitzer, 2014; Ni, Bouali-Benazzouz, Gao, Benabid, & Benazzouz, 2001; Ryu et al., 2011; Wichmann et al., 1999; Willard et al., 2019). The reasons for these inconsistencies are unclear but could relate to neuronal heterogeneity, homeostatic compensatory cellular and synaptic plasticity, species and technical differences in PD models, variation in recording techniques, and the influence of brain state and behavior on network dynamics. The therapeutic efficacy of high-frequency stimulation of the STN or GPi for the treatment of Parkinsonism further challenges the rate foundation of the classic model (Chiken & Nambu, 2016; McGregor & Nelson, 2019; Wichmann & DeLong, 2016).

In idiopathic and experimental PD, the GPe and STN exhibit an increase in correlated, phasic activity (Magill, Bolam, & Bevan, 2001; Mallet, Pogosyan, Marton, et al., 2008; Walters, Hu, Itoga, Parr-Brownlie, & Bergstrom, 2007). Thus, synchronous pauses in discharge and burst firing are relatively prevalent (Quiroga-Varela et al., 2013; Sanders, Clements, & Wichmann, 2013). In addition, these activity patterns can recur irregularly *or* rhythmically at frequencies between 1-35 Hz depending on the mode of dopamine depletion, species, behavior, and/or brain state (Delaville, McCoy, Gerber, Cruz, & Walters, 2015; Devergnas, Pittard, Bliwise, & Wichmann, 2014; Kuhn et al., 2009; Magill et al., 2001; Mallet, Pogosyan, Sharott, et al., 2008; Sharott, Vinciati, Nakamura, & Magill, 2017). Excessively correlated firing is thought to reduce the GPe-STN network’s capacity to encode information (Hammond et al., 2007; Mallet, Pogosyan, Sharott, et al., 2008), e.g., neuronal activity exhibits a marked reduction in somatotopic specificity (Bergman et al., 1994; Cho, Duke, Manzino, Sonsalla, & West, 2002; Ketzef et al., 2017; Leblois et al., 2006; Mallet, Pogosyan, Sharott, et al., 2008; Nambu, 2011; Pessiglione et al., 2005). The activity of the reciprocally connected GPe and STN can also become temporally offset, which given the convergence of these nuclei onto the GPi and SNr, may pathologically synchronize basal ganglia output (Bevan, Bolam, & Crossman, 1994; Moran et al., 2011; Shink, Bevan, Bolam, & Smith, 1996; Shouno, Tachibana, Nambu, & Doya, 2017; Tachibana, Iwamuro, Kita, Takada, & Nambu, 2011; Walters et al., 2007). The consistently therapeutic effect of manipulations that decorrelate network activity, e.g. STN deep brain stimulation or dopamine-based medications (Brown et al., 2001; Eusebio, Cagnan, & Brown, 2012; Heimer, Bar-Gad, Goldberg, & Bergman, 2002; Sharott et al., 2005; Whitmer et al., 2012), further support the linkage between excessively correlated GPe-STN network activity and motor dysfunction in PD. However, the origins of Parkinsonian GPe-STN network activity remain poorly understood. Hyperactivity (Escande, Taravini, Zold, Belforte, & Murer, 2016; H. Kita & Kita, 2011b; Mallet, Ballion, Le Moine, & Gonon, 2006; Moran et al., 2011; Sharott et al., 2017) and (mal)adaptive plasticity of D2-SPNs (Day et al., 2006; Fieblinger et al., 2014; Gittis et al., 2011; Taverna, Ilijic, & Surmeier, 2008), loss of autonomous GPe and STN activity (Chan et al., 2011; Hernandez et al., 2015; McIver et al., 2019; Shouno et al., 2017; C. L. Wilson et al., 2006; Zhu, Bartol, Shen, & Johnson, 2002), increases in the strength of GPe-STN inputs (Chu, Atherton, Wokosin, Surmeier, & Bevan, 2015; Fan, Baufreton, Surmeier, Chan, & Bevan, 2012; Moran et al., 2011), and/or downregulation of hyperdirect cortico-STN inputs (Chu, McIver, Kovaleski, Atherton, & Bevan, 2017; Mathai et al., 2015; Wang et al., 2018) have been hypothesized to play important roles. GPe and STN neurons are also considerably more heterogeneous than once thought, e.g. although the majority of GPe neurons project to the STN, “prototypic” GPe neurons also project to basal ganglia output neurons, midbrain dopamine neurons, and to the striatum where they preferentially target GABAergic interneurons (Bevan, Booth, Eaton, & Bolam, 1998; Mastro, Bouchard, Holt, & Gittis, 2014; Saunders, Huang, & Sabatini, 2016). In addition, “novel” classes of GPe neuron have been identified recently, including arkypallidal neurons, which comprise approximately one quarter of all GPe neurons and project only to the striatum (Abdi et al., 2015; Dodson et al., 2015; Hernandez et al., 2015; Mallet et al., 2012). Single- and sparse-cell labeling studies suggest that the targets of individual STN neurons are also more widespread and heterogeneous than originally thought (Hammond & Yelnik, 1983; H. Kita, Chang, & Kitai, 1983; Koshimizu, Fujiyama, Nakamura, Furuta, & Kaneko, 2013; Sato, Parent, Levesque, & Parent, 2000). While most STN neurons project to the GPi and SNr, some also innervate the GPe, the striatum, midbrain dopamine neurons, the brainstem, and the cortex (Degos et al., 2008; Hammond, Rouzaire-Dubois, Feger, Jackson, & Crossman, 1983; Jackson & Crossman, 1981; Hitoshi Kita & Kitai, 1987; Koshimizu et al., 2013; Parent & Hazrati, 1995; Sato et al., 2000; Smith, Hazrati, & Parent, 1990).

Previous studies in rats and mice utilized stereotyped cortical slow wave activity (SWA) and/or sensory stimulation evoked cortical activation (ACT) under anesthesia to probe the mechanisms underlying the abnormal cortical patterning of the dopamine-depleted basal ganglia in the absence of confounding behavioral variables like sensorimotor feedback (Magill et al., 2001; Mallet, Pogosyan, Marton, et al., 2008; Mallet, Pogosyan, Sharott, et al., 2008; Walters et al., 2007; Zold, Escande, Pomata, Riquelme, & Murer, 2012). Here, we adopted a similar approach but combined it with *in vivo* optogenetic inhibition to selectively identify and suppress the activity of D2-SPNs, parvalbumin (PV)-expressing prototypic GPe neurons, and STN neurons to directly assess their contributions to GPe-STN network activity in both dopamine-intact and -depleted mice. In addition, we compared the autonomous activity of PV-expressing prototypic GPe neurons in brain slices from control and Parkinsonian mice. Through these approaches, we elucidate the contribution of specific circuit elements to the patterning of GPe-STN network activity in normal and Parkinsonian mice.

## Results

Data are presented as median and interquartile range (IQR). Paired and unpaired data were compared using the non-parametric Mann-Whitney U (MWU) and Wilcoxon Signed Rank (WSR) tests, respectively, and Fisher’s exact test was used for contingency analyses. When applicable, the Holm-Bonferroni correction for multiple comparisons (Holm, 1997) was applied (adjusted p-values are indicated as p_h#_, where # is the adjustment factor). P values <0.05 were considered significant.

To determine the activity of D2-SPNs, prototypic PV-expressing GPe neurons, and STN neurons and their impact on GPe-STN network activity, neurons were identified and their activity manipulated through activation of the inhibitory opsin Arch (Chow et al., 2010). Arch-GFP was expressed in D2-SPNs, prototypic PV-expressing GPe neurons, and STN neurons through injection of an AAV vector carrying a cre-dependent expression construct in the striatum of A2A-cre mice, the GPe of PV-cre mice, and the STN of GABRR3-cre mice, respectively. During the same surgery, mice also received a unilateral injection of vehicle or 6-hydroxydopamine (6-OHDA) in the medial forebrain bundle (MFB) as a surgical control and to produce degeneration of midbrain dopamine neurons, respectively. 18.5, 15-21 days after surgery mice were anesthetized with a combination of urethane and ketamine/xylazine and striatal, GPe, and STN neuronal activity in the ipsilateral hemisphere was recorded and/or manipulated using silicon tetrodes/optrodes. Coincident cortical activity was assessed from the electroencephalogram (EEG) that was obtained from a peridural screw “electrode” affixed over the primary motor cortex. The relative powers of the EEG in the 0.5-1.5 Hz, 10-39.9 Hz, and 40-250 Hz bands were not significantly different in vehicle- and 6-OHDA-injected mice (Table S1). Dopaminergic innervation of the striatum in 6-OHDA-injected mice and vehicle-injected mice was assessed through immunohistochemistry for tyrosine hydroxylase (TH) (Table S2).

### In dopamine-depleted mice D2-SPNs were hyperactive during cortical SWA

The specificity of Arch-GFP expression in D2-SPNs in A2A-cre mice was confirmed by its presence in the subset of SPNs that project to the GPe (Fig. 1A-D) but not to the SNr (Fig. 1E-F). In regions expressing Arch-GFP, approximately one half of striatal neurons were directly inhibited by activation of Arch-GFP and therefore identified as D2-SPNs (vehicle: 55 %, n = 62 of 112; 6-OHDA: 56 %, n = 35 of 63; p = 0.2076; Fisher’s Exact). Of the neurons that were excluded due to the absence of direct inhibition, 2 % (n = 1 of 50) in vehicle- and 12 % (n = 3 of 25) in 6-OHDA-treated animals were disinhibited during optogenetic stimulation. In both vehicle- and 6-OHDA-injected animals, D2-SPN activity was phasic and entrained to the active (positive) component of cortical SWA in the EEG (Fig. 2A, B). Because dopamine negatively modulates both cortico-striatal transmission (Bamford et al., 2004; Higley & Sabatini, 2010) and the intrinsic excitability of D2-SPNs (Gerfen & Surmeier, 2011; Planert, Berger, & Silberberg, 2013), loss of dopamine was predicted to increase cortical SWA-associated D2-SPN activity. Consistent with this prediction, the frequency and the coefficient of variation of the interspike interval (CV) of D2-SPN firing measured during 30 seconds epochs of cortical SWA were elevated in 6-OHDA-injected mice relative to vehicle controls (Fig. 2A-C; vehicle: rate = 1.75, 0.918-3.55 Hz; n = 62; 6-OHDA: rate = 3.02, 1.53-4.58 Hz; n = 34; p = 0.02068; MWU; vehicle: CV = 1.47, 1.19-1.77; n = 62; 6-OHDA: CV = 1.646, 1.25-2.179; n = 34; p = 0.04235; MWU). The relationship between cortical SWA and D2-SPN firing was further examined through the generation of phase histograms. First, each spike was assigned to the phase of the EEG, with 0°/360° and 180° corresponding to the peak active and inactive components of cortical SWA, respectively. Data were binned at 10° in figures, and at 180° for statistical comparisons. Each neuron’s spike probabilities were calculated from (spikes/bin)/(total # of spikes) X 100%. Firing in-phase to cortical SWA was designated as activity within a 180° window centered on 0/360°. Anti-phase firing was designated as activity within a 180° window centered on 180°. The ratio of in-:anti-phase spike probability in D2-SPNs was not altered by dopamine depletion (Fig. 2C, D; vehicle = 9.814, 3.60-14.4; n = 48; 6-OHDA = 3.91, 2.51-17; n = 27; p = 0.4671; MWU). Together these data demonstrate that following dopamine depletion, D2-SPNs are hyperactive during cortical SWA but their phase relationship to cortical SWA is unaltered. These data in mice are consistent with the activity of putative and identified striatopallidal neuron activity in dopamine intact and - depleted rats during cortical SWA (Escande et al., 2016; Mallet et al., 2006; Sharott et al., 2017; Walters et al., 2007; Zold et al., 2012).

**Figure 1.**
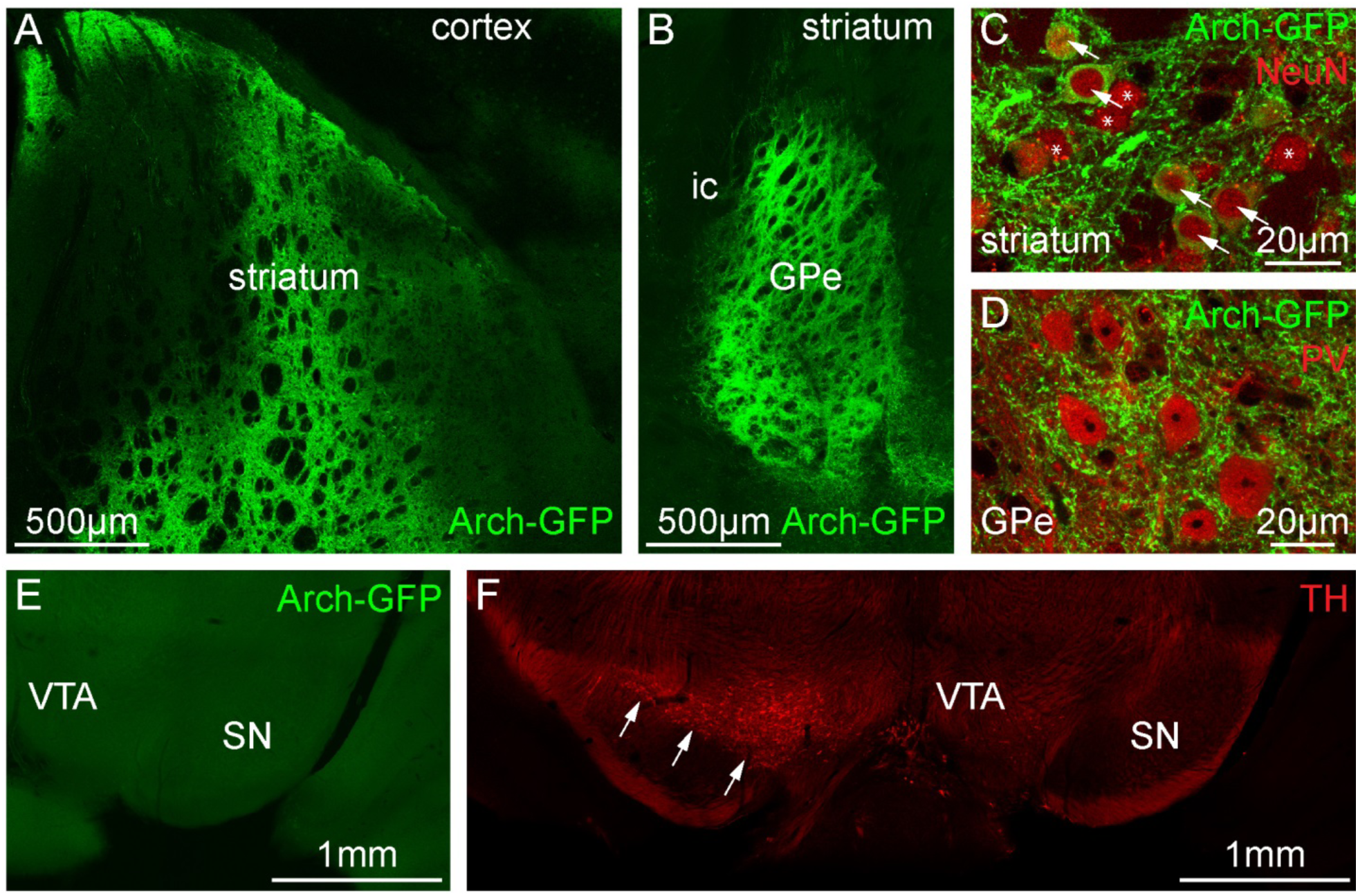
Cre-dependent viral expression of Arch-GFP in D2-SPNs in A2A-cre mice. (A-D) Expression of Arch-GFP (green) in D2-SPNs in the striatum (A, C) and their axon terminals in the GPe (B, D; ic, internal capsule). (C, D) Expression of NeuN (red) in the somata of striatal (C) and GPe (D) neurons. (C) Arch-GFP-expressing and non-expressing NeuN-immunoreactive SPNs are denoted by arrows and asterisks, respectively. (E, F) Absence of Arch-GFP expressing axon terminals (E) and TH-expressing neurons (F) ipsilateral to injections of AAV in the striatum (A) and 6-OHDA in the MFB. TH-immunoreactive substantia nigra (SN) and ventral tegmental area (VTA) neurons (red; arrows) are visible contralateral to the injection of 6-OHDA.

**Figure 2.**
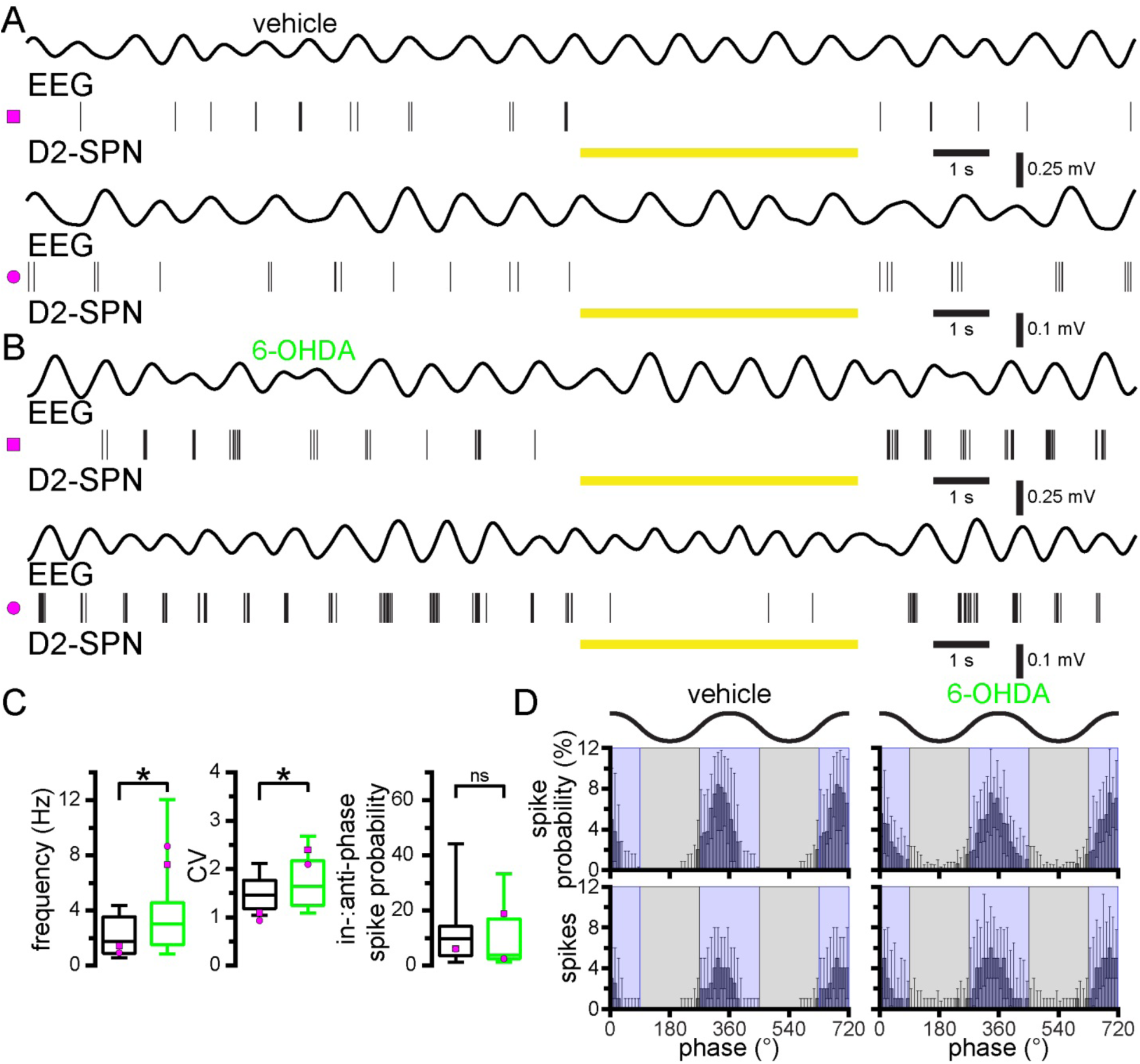
D2-SPNs are hyperactive in dopamine-depleted mice. (A-B) Representative EEG (upper trace; filtered at 0.5-1.5 Hz) and D2-SPN unit activity (below) from vehicle-(A) and 6-OHDA-injected (B) mice. D2-SPNs were identified by their inhibited activity during optogenetic stimulation of Arch-GFP (yellow bar). (C-D) The frequency and CV of D2-SPN activity were significantly greater in 6-OHDA-versus vehicle-injected mice but D2-SPN activity was similarly entrained to the active component of SWA. (C) Population box plots, D2-SPN firing rate (left), CV of the ISI (middle), and in-:anti-phase spike probability (right); representative examples plotted with magenta symbols. (D) Population linear phase histograms illustrate the phase relationship between D2-SPN neuron firing and SWA in vehicle- and 6-OHDA-injected mice. Active (blue) and inactive (gray) components of SWA are denoted. *, p < 0.05. ns, not significant.

### In dopamine-depleted mice prototypic PV GPe neuron activity was relatively anti-phasic to cortical SWA

To identify prototypic GPe neurons, the majority of which express PV, Arch-GFP was expressed in a cre-dependent manner in PV-cre mice (Abdi et al., 2015; Dodson et al., 2015; Hernandez et al., 2015; Mastro et al., 2014). Immunohistochemistry confirmed that the majority of Arch-GFP-expressing neurons co-expressed PV (Fig. 3A, B; 83 % of Arch-GFP-expressing neurons co-expressed PV; 73-93 %; N = 2 mice). Consistent with their prototypic identity, Arch-GFP was also strongly expressed by their axon terminals in the STN (Fig. 3C, D). Incomplete immunohistochemical detection of PV was presumably the reason for the absence of PV immunoreactivity in 17 % of Arch-GFP expressing GPe neurons. In contrast, arkypallidal neurons, identified by their immunoreactivity for FoxP2 (Abdi et al., 2015; Dodson et al., 2015; Hernandez et al., 2015), did not co-express Arch-GFP (Fig. 3B; 0 %; N = 2 mice). Together these data confirm the selective expression of Arch-GFP in prototypic PV GPe neurons and the absence of Arch-GFP expression in arkypallidal FoxP2 GPe neurons. Consistent with the relative abundance of prototypic PV GPe neurons, the majority of GPe neurons that were recorded were inhibited by optogenetic activation of Arch-GFP (vehicle: 82 %, n = 27 of 33; 6-OHDA: 84 %, n = 27 of 32; p = 1.0000; Fisher’s Exact). However, given that recordings were initiated only from tetrode locations where at least one GPe neuron was inhibited during stimulation, recordings were likely to be biased toward prototypic neurons. Of those neurons that were not directly inhibited by stimulation, 83 % (n = 5 of 6) in vehicle and 40 % (n = 2 of 5) in 6-OHDA-treated mice were disinhibited, presumably due to inhibition of local collaterals.

**Figure 3.**
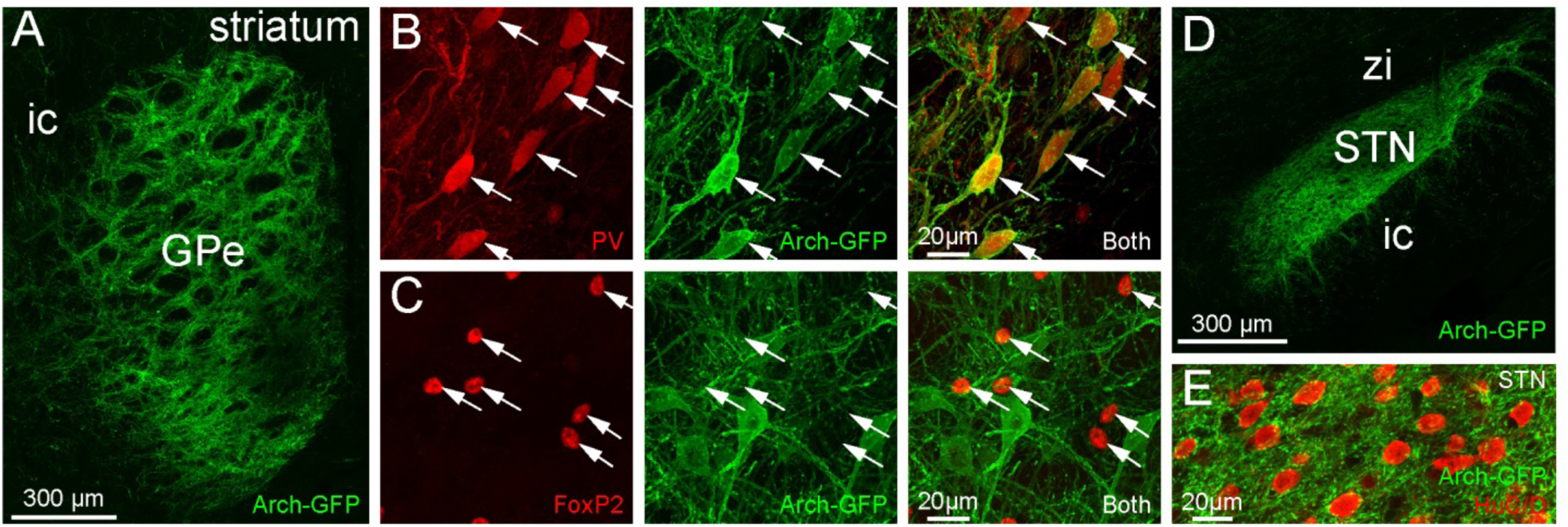
Cre-dependent viral expression of Arch-GFP in PV GPe neurons in PV-cre mice. (A-E) Expression of Arch-GFP (green) in PV GPe neurons (A-C) and their axon terminals in the STN (D, E; zi, zona incerta). (A-C) Expression of Arch-GFP in the GPe. (B) Expression of Arch-GFP in PV-immunoreactive prototypic GPe neurons (red; arrows). (C) Absence of Arch-GFP expression in FoxP2-immunoreactive arkypallidal GPe neurons (red; arrows). (D) Arch-GFP expression in GPe axon terminals in the STN arising from the injection in A. (E) Arch-GFP expressing axon terminals in the vicinity of NeuN-immunoreactive STN neurons (red).

As for D2-SPNs, PV GPe neuron firing was first recorded during 30 second epochs of robust cortical SWA. In control mice, PV GPe neurons discharged in a tonic, irregular firing pattern during cortical SWA (Fig. 4A, C; vehicle: frequency = 16.8, 9.23-30.1 Hz; CV = 0.868, 0.554-1.01; n = 27). In dopamine-depleted mice, the rate of discharge of PV GPe neurons was similar but firing was more irregular relative to control (Fig. 4B, C; 6-OHDA: frequency = 15.6, 12.5-25.2 Hz; n = 27; p = 0.9280; MWU; CV = 1.191, 0.824-1.59; n = 27; p = 2.163e-3; MWU). Comparison of the in-:anti-phase spike probability ratio in vehicle- and 6-OHDA injected mice, revealed a shift in PV GPe neuron phase preference to firing that was more anti-phasic to cortical SWA (Fig. 4C, D; SWA: vehicle = 0.801, 0.664-1.18; n = 27; 6-OHDA = 0.675, 0.418-0.883; n = 27; p = 9.160e-3; MWU). These data are consistent with studies in dopamine-depleted rats, which demonstrated that juxtacellularly labeled prototypic GPe neurons, defined on the basis of their descending axon collaterals and in some cases their expression of PV, exhibited firing that was relatively anti-phasic to cortical SWA (Abdi et al., 2015; Magill et al., 2001; Mallet et al., 2012; Mallet, Pogosyan, Sharott, et al., 2008). Past studies focused on prototypic GPe neuron populations that adhered to specific firing characteristics such as phase-locking to SWA for analysis. Here, we included all optogenetically identified PV GPe neurons and report more variability in phase preference. Our use of mice versus rats in earlier studies may also contribute to this variability.

**Figure 4.**
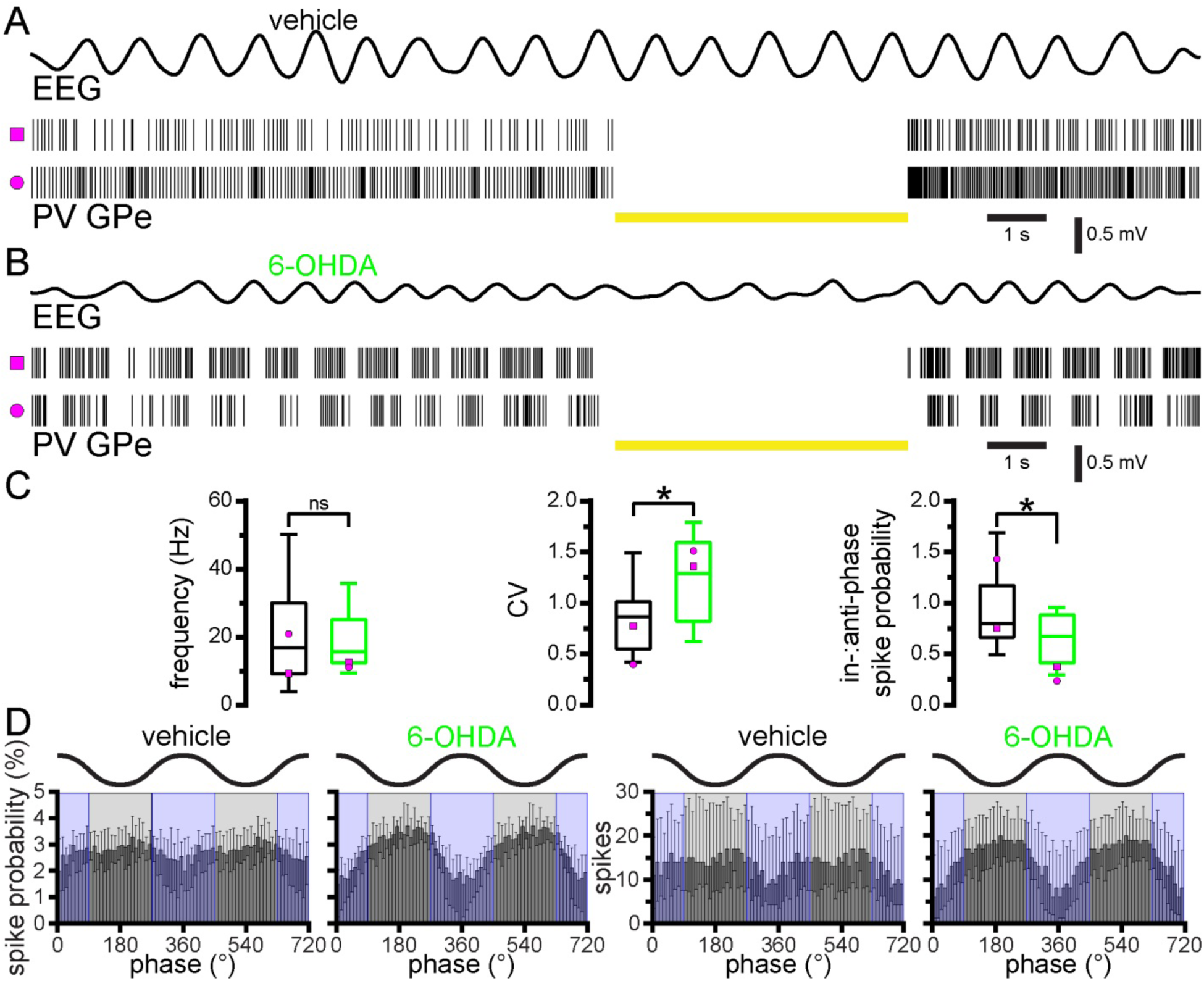
Prototypic PV GPe neuron activity is relatively anti-phasic to cortical SWA in 6-OHDA-injected mice. (A, C, D) In vehicle-injected dopamine-intact mice PV GPe neuron activity was relatively tonic and not consistently phase-related to SWA. (B-D) In 6-OHDA-injected, dopamine-depleted mice PV GPe neuron activity was relatively entrained to the inactive component of cortical SWA. (A, B) PV GPe neurons were identified through optogenetic stimulation of Arch-GFP which inhibited their activity (yellow bar). (A, B) Representative examples of EEG and associated PV GPe neuron activity. (C) Population data, PV GPe neuron firing rate (left), CV of the ISI (middle), and in-:anti-phase spike probability (right); example data plotted with magenta symbols. (D) Population linear phase histograms of PV GPe neuron firing relative to cortical SWA in vehicle- and 6-OHDA-injected mice. *, p < 0.05. ns, not significant.

### In dopamine-depleted mice optogenetic inhibition of hyperactive D2-SPNs reduced anti-phasic GPe firing

To determine whether the increase in anti-phasic GPe activity in dopamine-depleted mice is linked to the hyperactivity of D2-SPNs, D2-SPNs were optogenetically inhibited for 5 second epochs and the effect on GPe neuron activity was assessed (Table S1). Use of A2A-cre mice to selectively express Arch-GFP in D2-SPNs precluded optogenetic identification of prototypic PV GPe neurons. Although recordings were only initiated where putative disinhibition of GPe neurons was observed during optical stimulation, responsive GPe neurons were relatively rare (46 %, n = 26 of 57), presumably because 1) the zone of optogenetic inhibition versus the size of the striatum was small and 2) the striatopallidal projection is highly topographic in nature (Chang, Wilson, & Kitai, 1981; Hazrati & Parent, 1992; Hedreen & DeLong, 1991; Smith, Bevan, Shink, & Bolam, 1998; C. J. Wilson & Phelan, 1982), lowering the probability of recording from connected parts of the striatum and GPe (Fig. 5A, B). In the majority of responsive GPe neurons D2-SPN inhibition elevated the frequency (Fig. 5A, B; 6-OHDA: 85 %, n = 22 of 26; laser off = 17.9, 11.3-29.4 Hz; laser on = 23.0, 15.9-34.0 Hz; n = 26; p = 1.013e-6; WSR) and regularity (Fig. 5A, B; 6-OHDA: 92 %, n = 23 of 25; laser off CV = 1.10, 0.898-1.27; laser on CV = 0.701, 0.561-0.817; n = 25; p = 2.98e-7; WSR) of their firing. Furthermore, inhibition of D2-SPNs reduced anti-phasic GPe activity, as indicated by an increase in in-:anti-phase spike probability (Fig. 5B, C; 6-OHDA: 64 %, n = 16 of 25; laser off = 0.694, 0.567-1.07; laser on = 0.897, 0.791-1.02; n = 25; p = 0.04826; WSR).

**Figure 5.**
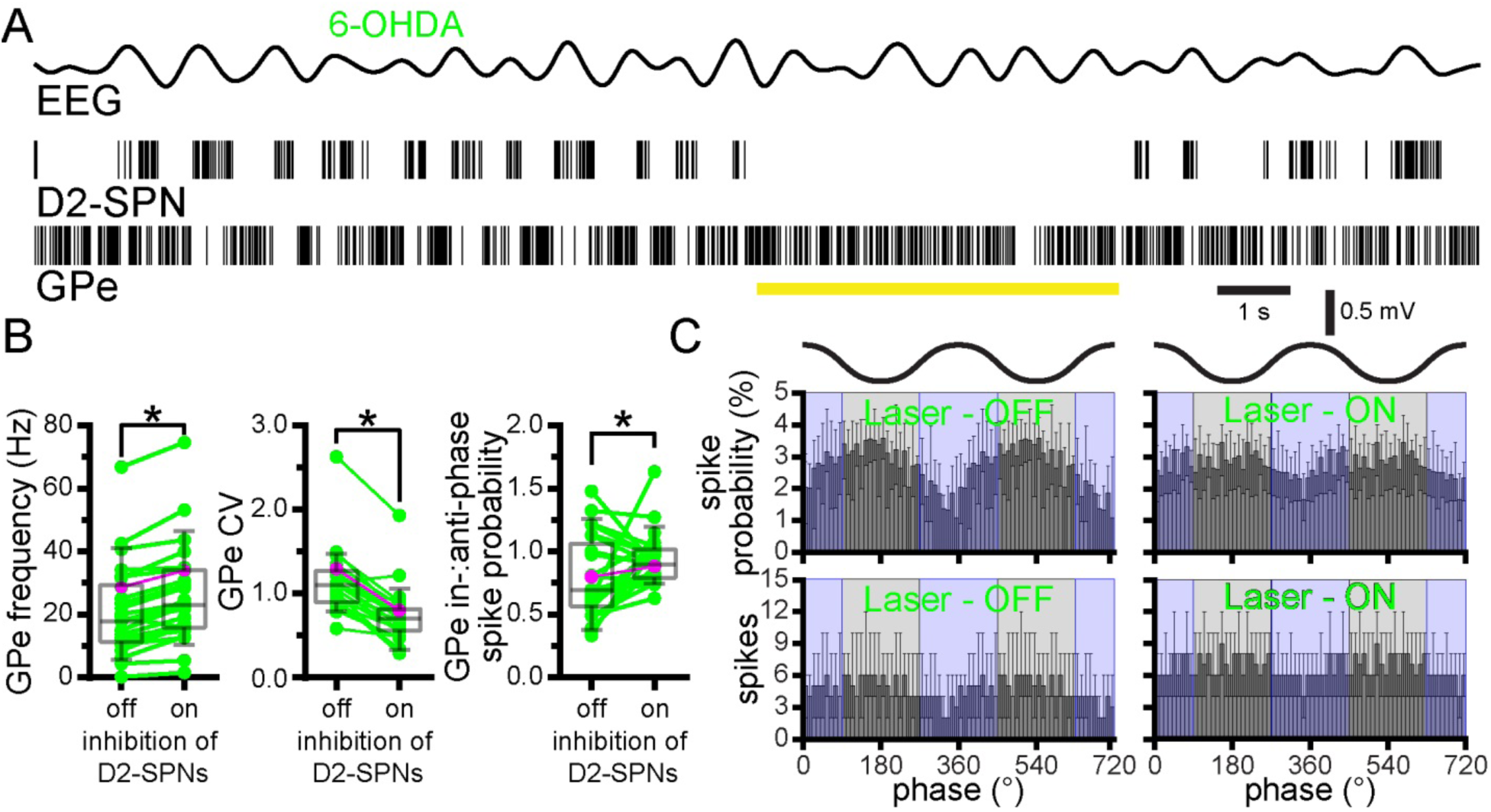
Optogenetic inhibition of D2-SPNs reduced anti-phasic GPe activity in 6-OHDA-injected mice. (A-C) During SWA optogenetic inhibition of D2-SPNs (yellow bar; laser on) in 6-OHDA-injected mice increased the frequency and decreased the variability (CV) of firing, and increased the in-:anti-phase spike probability of unidentified GPe neurons (A, representative example; B, population data, firing rate (left), CV of the ISI (middle), and in-:anti-phase spike probability (right); example data plotted in magenta; C, population linear phase histograms). *, p < 0.05.

To more selectively examine the impact of D2-SPN input on prototypic PV GPe neurons, putative PV GPe neuron activity was isolated by analyzing neurons with in-:anti-phase spike probability ratios falling within the IQR of identified PV GPe neurons in PV-cre mice (PV-like GPe neurons). In response to optogenetic inhibition of D2-SPNs, PV-like GPe neurons in A2A-cre mice were uniformly disinhibited, (Fig. 5-1 A; 6-OHDA: 100 %, n = 14 of 14; laser off = 24.3, 16.3-34.2Hz; laser on = 30, 20.5-40.9Hz; n = 14; p = 1.221e-4; WSR). In addition, their firing decreased in irregularity (Fig. 5-1A; 6-OHDA: 93 %, n = 13 of 14; laser off CV = 1.24, 0.967-1.37; laser on CV = 0.752, 0.536-0.870; n = 14; p = 2.441e-4; WSR) and became less anti-phasic (Fig. 5-1A, B; 6-OHDA: 93 %, n = 13 of 14; laser off = 0.638, 0.574-721; laser on = 0.813, 0.76-102; n = 14; p = 2.441e-4; WSR). Thus, optogenetic inhibition of D2-SPNs reduced anti-phasic discharge and normalized the pattern of unidentified GPe neuron activity in dopamine-depleted mice. These effects were more uniform in the PV-like subset of GPe neurons. Together, these data demonstrate that during cortical SWA, hyperactive D2-SPNs contribute to the abnormal anti-phasic firing of GPe neurons in dopamine-depleted mice.

**Figure 5-1.**
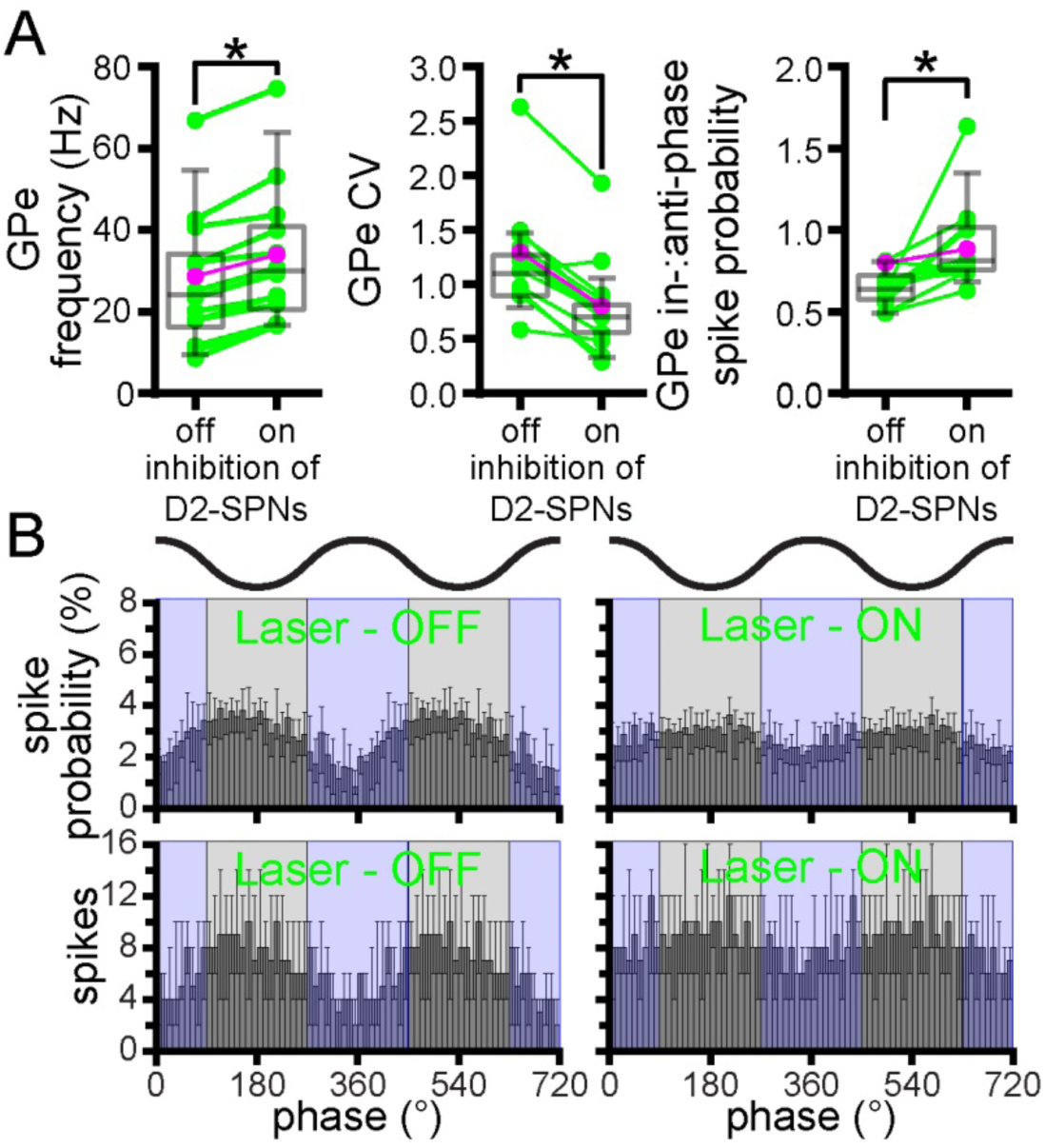
Putative PV GPe neurons are relatively uniform in their response to D2-SPN inhibition in 6-OHDA-injected mice. (A-B) Similar to the unidentified population of GPe neurons, optogenetic inhibition of D2-SPNs in 6-OHDA-injected mice increased the frequency and decreased the variability (CV) of firing, and increased the in-:anti-phase spike probability in putative PV GPe neurons (A, representative examples; B, population data, firing rate (left), CV (middle), and in-:anti-phase spike probability (right); example cell data from Fig. 5 plotted in magenta; C, population linear phase histograms). *, p < 0.05.

### The autonomous firing of PV GPe neurons is significantly elevated following loss of dopamine

Loss of autonomous activity in GPe neurons has been suggested to contribute to their abnormal activity *in vivo* following dopamine depletion (Chan et al., 2011). However, this work was carried out before the recent discovery of multiple GPe neuron subtypes. Therefore, the autonomous firing of prototypic PV GPe neurons was compared in *ex vivo* brain slices derived from vehicle- and 6-OHDA-injected mice (Table S1). In order to identify PV GPe neurons for patch clamp recording, tdTomato was conditionally expressed in PV neurons by crossing PV-cre mice with Ai9 reporter mice in which a loxP-flanked STOP cassette prevents transcription of downstream tdTomato. Selective expression of tdTomato in PV GPe neurons was first confirmed using immunohistochemistry for PV. Of neurons expressing tdtomato, PV, or both markers: 1) 73.2, 66.1-77.9 % (Fig. 6A-E; n = 3 mice) co-expressed tdTomato and PV 2) 8.9, 4.1-14.2 % (Fig. 6A-E; n = 3 mice) expressed tdtomato but were not immunoreactive for PV, most likely due to less than 100% efficiency of immunodetection 3) 18.0, 18.0-19.8 % (Fig. 6A-E; n = 3 mice) of neurons did not express tdTomato but were PV-immunoreactive, presumably due to < 100 % efficiency of cre-mediated excision of the loxP-flanked STOP cassette. Less than 1 % of tdTomato expressing GPe neurons co-expressed the arkypallidal neuron marker FoxP2, consistent with previous reports (Abdi et al., 2015; Hernandez et al., 2015; data not shown). Together, these data suggest that tdTomato expression in this mouse line is a reliable marker of prototypic, PV GPe neurons. Cell-attached, current clamp recordings of GPe tdTomato-expressing neurons in *ex vivo* brain slices were conducted in the presence of AMPA, NMDA, GABA_A_, and GABA_B_ receptor antagonists in order to measure their autonomous activity. As described previously, PV GPe neurons discharged regularly and at high frequency in brain slices derived from dopamine-intact mice (Fig. 6F, G; vehicle: frequency = 35.1, 26.8-44.0 Hz; CV = 0.13, 0.10-0.23; n = 89) (Abdi et al., 2015; Dodson et al., 2015; Hernandez et al., 2015; Mastro et al., 2014). The autonomous firing of PV GPe neurons in slices from 6-OHDA-injected mice was not only retained but significantly elevated compared to firing of PV GPe neurons in vehicle-injected mice (Fig. 6F, G; 6-OHDA: frequency = 41.7, 34.0-50.2 Hz; CV = 0.12, 0.08-0.20; n = 98; frequency, p = 5.35e-4; CV, p = 0.1336; WSR). Together these data suggest that the abnormally phasic pattern of PV GPe neuron firing in dopamine-depleted mice *in vivo* is not caused by loss of autonomous firing. In fact, autonomous firing was significantly up-regulated following the loss of dopamine, presumably through engagement of homeostatic compensatory mechanisms that were triggered by an increase in striatopallidal transmission.

**Figure 6.**
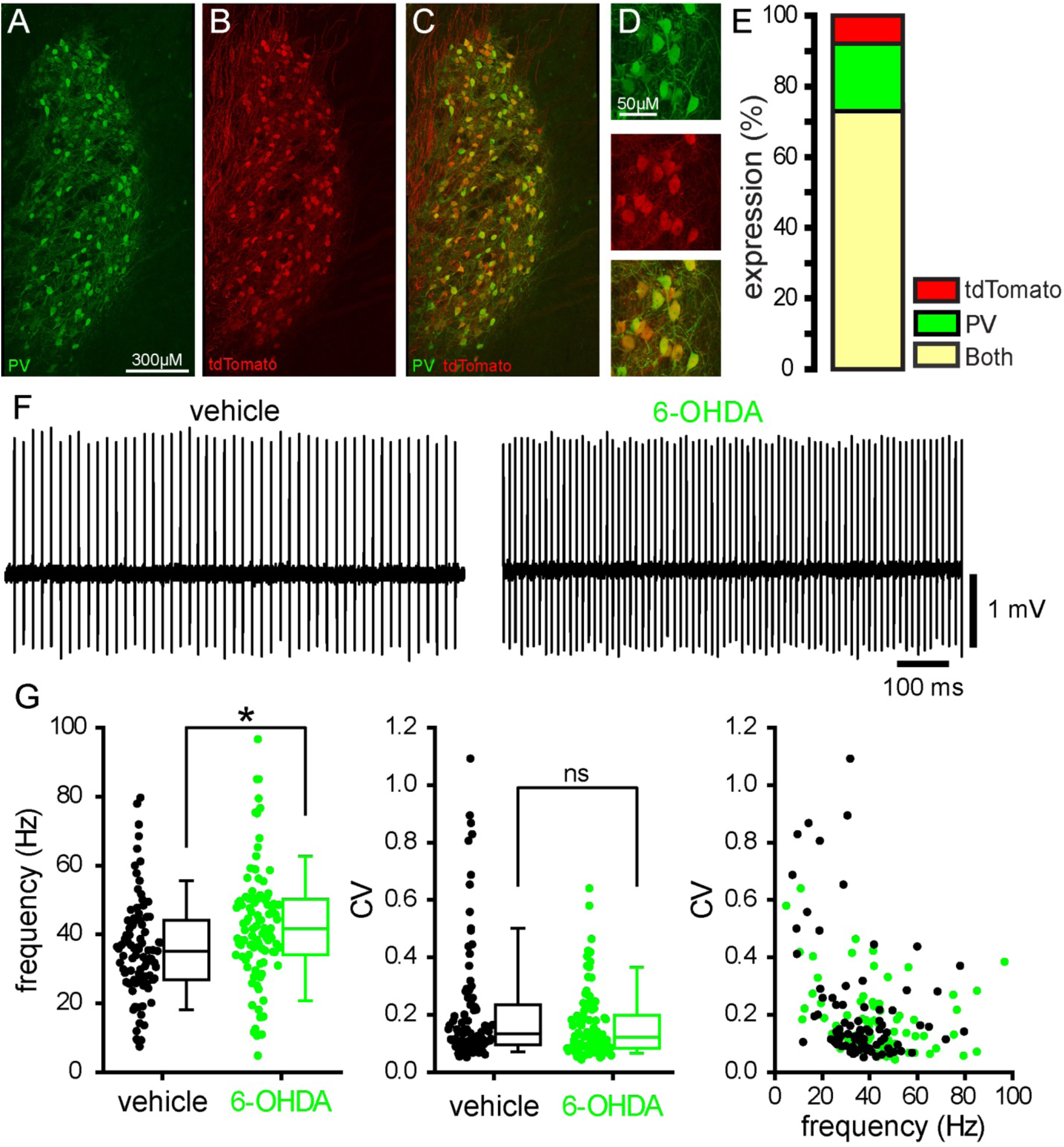
The frequency of autonomous prototypic PV GPe neuron activity *ex vivo* is elevated in 6-OHDA-injected mice. (A-E) In PV-cre X Ai9 mice the majority of GPe neurons that expressed PV (green) or tdTomato (red) expressed both proteins (yellow), as evinced by a representative coronal section through the GPe (A-D; D, upper panel, PV; middle panel, tdTomato, lower panel, both) and quantitative population data (E). (F-G) The autonomous activity of tdTomato-expressing GPe neurons recorded in the loose-seal cell-attached configuration was not disrupted in 6-OHDA-injected mice relative to activity in control mice. Indeed, the frequency of autonomous firing was significantly greater in 6-OHDA-injected dopamine-depleted mice compared to that in control mice. (F) Representative examples. (G) Population data. *, p < 0.05. ns, not significant.

### Optogenetic inhibition of STN neurons reduced GPe neuron activity in dopamine-intact and -depleted mice and increased anti-phasic GPe neuron activity in dopamine-depleted mice

Reciprocally connected GPe-STN neurons have been proposed as a generator of abnormal oscillatory activity in PD (Holgado, Terry, & Bogacz, 2010; Moran et al., 2011; Plenz & Kital, 1999). To determine the role of the STN in patterning GPe activity, we compared the responses of GPe neurons to optogenetic inhibition of STN neurons for 5 seconds in vehicle- and 6-OHDA-injected GABRR3-cre mice during SWA (Table S1). Confocal imaging confirmed the robust and selective expression of Arch-GFP in STN neurons (Fig. 7A, C) and their axon terminals in the GPe (Fig. 7B, D). However, the use of GABRR3-cre mice to selectively express Arch-GFP in the STN precluded optogenetic identification of prototypic PV GPe neurons. The proportion of GPe neurons that responded to optogenetic inhibition of the STN was not significantly different in vehicle- and 6-OHDA-injected mice (vehicle: 62 % responsive, n = 23 of 37; 6-OHDA 70 % responsive, n = 21 of 30; p = 0.6076; Fisher’s Exact). Optogenetic inhibition of STN neurons reduced GPe neuron activity in all responsive neurons in both vehicle- and 6-OHDA-injected mice (Fig. 8 A-C; vehicle: laser off = 14.3, 7.60-21.5 Hz; laser on = 7.90, 4.90-12.6 Hz; n = 23; p_h4_ = 9.54e-7; WSR; 6-OHDA: laser off = 17.8, 13.1-28.3 Hz; laser on = 15.6, 7.50-23.2 Hz; n = 21; p_h3_ = 8.58e-6; WSR). The firing rate of GPe neurons in vehicle-injected mice was not significantly different to those in 6-OHDA-injected mice (Fig. 8 A-C; laser off: p_h1_ = 0.05722; MWU); however, during optogenetic inhibition of the STN, GPe neuron firing rates were relatively elevated in 6-OHDA mice (Fig. 8 A-C; laser on: p_h2_ = 0.02696; MWU).

**Figure 7.**
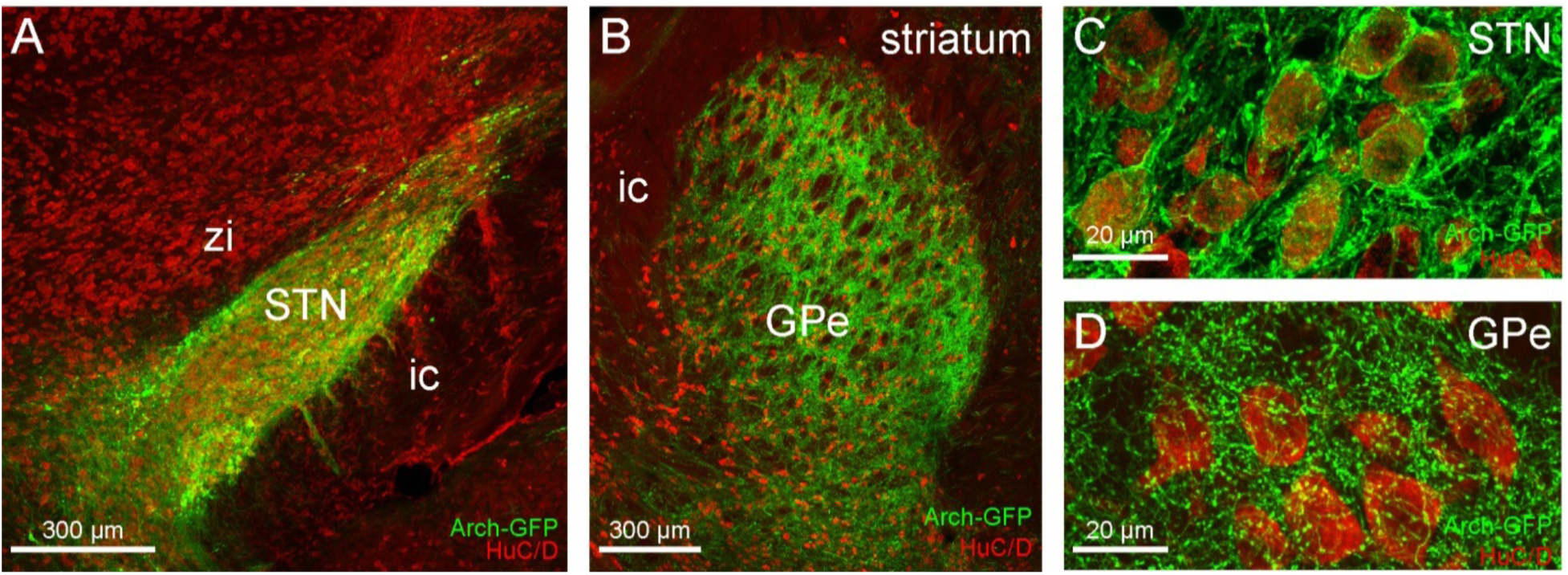
Cre-dependent viral expression of Arch-GFP in the STN of GABRR3-cre mice. (A-D) Coronal sections illustrating expression of Arch-GFP (green) in STN neuron somata (A, C) and their axon terminals in the GPe (B, D) in GABRR3-cre mice. (A, C) Expression of Arch-GFP in the STN. (C) Expression of Arch-GFP in NeuN-immunoreactive (red) STN neurons. (B, D) Arch-GFP expression in the GPe arising from the injection in A. (D) Arch-GFP expressing STN axon terminals in the vicinity of NeuN-immunoreactive (red) GPe neurons.

**Figure 8.**
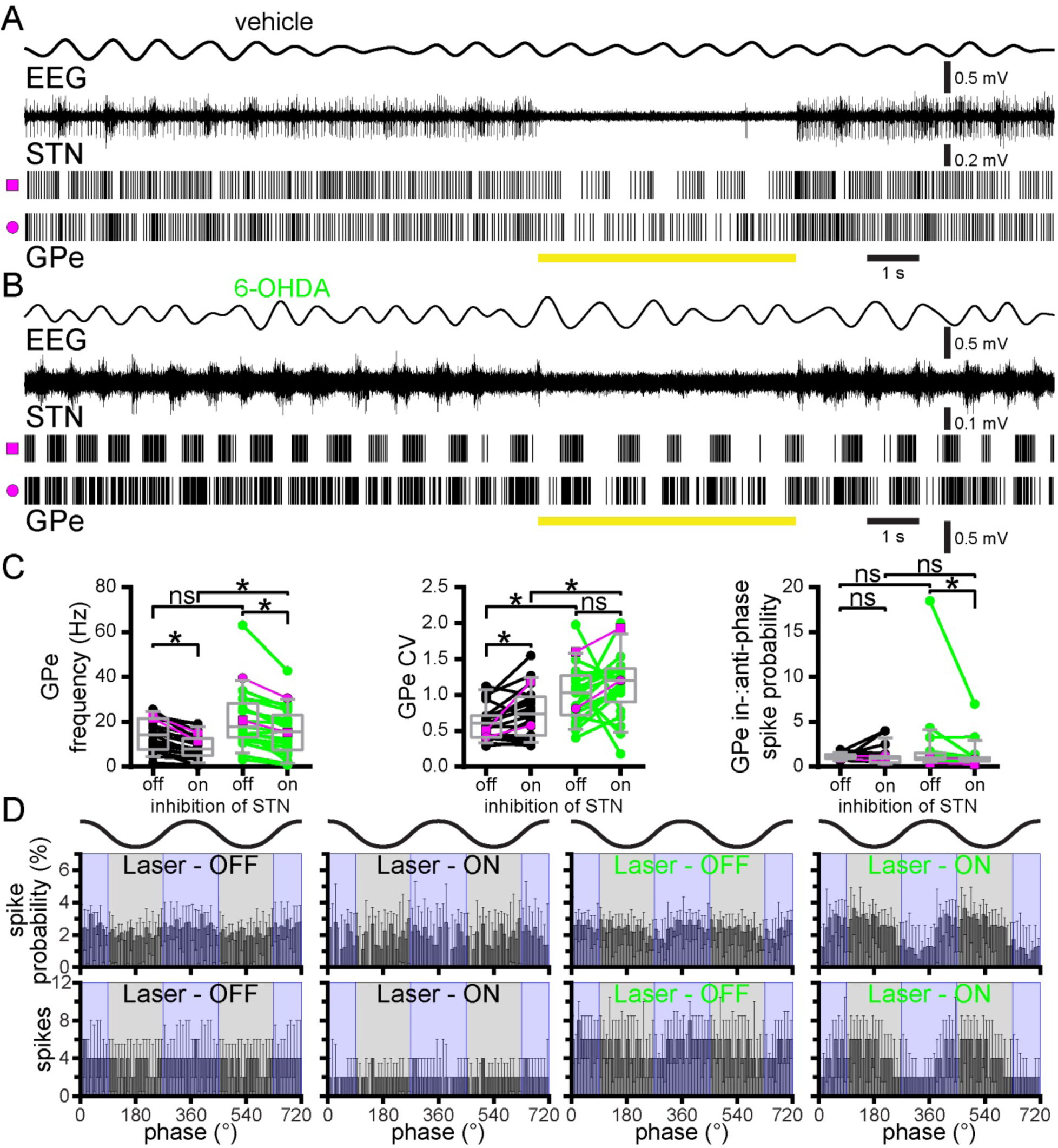
In 6-OHDA-injected mice optogenetic inhibition of STN neurons decreased the firing rate of unidentified GPe neurons but increased their anti-phasic activity. (A-D) Optogenetic inhibition of STN activity in vehicle- and 6-OHDA-injected GABRR3-cre mice reduced the frequency of unidentified GPe neuron firing. In 6-OHDA-injected mice inhibition of the STN also significantly reduced the in-:anti-phase spike probability of GPe neurons. (A, B) Representative examples. Note multi-unit STN activity is illustrated due to the difficulty of spike sorting individual STN units when they were recorded with an optrode compared to silicon tetrodes without a fiber optic. (C) Population data, firing rate (left), CV (middle), and in-:anti-phase spike probability (right); example data plotted in magenta. (D) Linear phase histograms of GPe neuron activity prior to and during optogenetic inhibition of STN neurons. *, p < 0.05. ns, not significant.

The firing of unidentified GPe neurons in vehicle- and 6-OHDA-injected mice prior to and during optogenetic inhibition of the STN was significantly more irregular in 6-OHDA-injected mice (Fig. 8 A-C; vehicle: laser off CV = 0.606, 0.413-0.717, n = 23; 6-OHDA: laser off CV = 1.03, 0.723-1.27, n = 21; p_h4_ = 2.502e-3; MWU; vehicle: laser on CV = 0.737, 0.436-0.973; 6-OHDA: laser on CV = 1.20, 0.903-1.37; n = 21; p_h3_ = 6.951e-3; MWU). During optogenetic inhibition of STN neurons, the regularity of firing of responsive GPe neurons decreased in vehicle-injected mice but was not altered in 6-OHDA-injected mice (Fig. 8 A-C; vehicle: laser off CV = 0.581, 0.410-0.711, n = 22; vehicle: laser on CV = 0.737, 0.436-0.973; n = 22; p_h2_ = 9.37e-3; WSR; 6-OHDA: laser off CV = 1.03, 0.723-1.27, n = 21; 6-OHDA: laser on CV = 1.20, 0.903-1.37; n = 21; p_h1_ = 0.1281; WSR). During optogenetic inhibition of the STN, the in-:anti-phase spike probability ratio of GPe neurons decreased in 6-OHDA- but was unaltered in vehicle-injected mice (Fig. 8 A-C; vehicle: laser off = 1.18, 0.908-1.36; laser on = 1.11, 0.571-1.43; n = 22; p_h2_ = 1.000; WSR; 6-OHDA: laser off = 1.02, 0.799-1.55; laser on = 0.781, 0.641-1.02; n = 21; p_h4_ = 0.01143; WSR). Unidentified GPe neurons were less anti-phasic than identified PV GPe neurons in dopamine-depleted mice. The reason for this difference is likely due to the inclusion of arkypallidal GPe neurons thought to fire in-phase with SWA following dopamine depletion and perhaps also PV-prototypic GPe neurons, whose phase preference has not been well characterized (Abdi et al., 2015; Mallet et al., 2012). It is also possible that during SWA pauses in prototypic GPe neuron firing in 6-OHDA-injected mice enhance the firing of arkypallidal neurons through disinhibition, thus increasing in-phase firing in the unidentified GPe neuron population.

To examine the impact of STN inhibition on PV GPe neuron firing, PV-like GPe neurons in 6-OHDA-injected mice were isolated from the unidentified population as for A2A-cre mice on the basis of their in-:anti-phase spike probability. The firing rate of all PV-like GPe neurons decreased during optogenetic inhibition of the STN (Fig.8-1 A, B; 6-OHDA: laser off = 26.2, 20.3-33.9 Hz; laser on = 19.4, 15.3-27.9 Hz; n = 7; p = 0.01562; WSR). Unlike the unidentified population of GPe neurons, the irregularity of PV-like GPe neurons was elevated by optogenetic inhibition of the STN (Fig.8-1 A, B; 6-OHDA: laser off CV = 0.815, 0.509-1.3 Hz; laser on CV = 1.17, 1.03-1.93; n = 7; p = 0.01562; WSR). Lastly, all PV-like GPe neurons became more anti-phasic during optogenetic inhibition of the STN as represented by a significant decrease in in-:anti-phase spike probability (Fig.8-1 A, B; 6-OHDA: laser off = 0.795, 0.670-0811 Hz; laser on = 0.731, 0.28-0.781; n = 7; p = 0.01562; WSR). Overall, STN silencing enhanced anti-phasic firing in unidentified and PV-like GPe neurons, arguing that STN-GPe transmission opposes rather than facilitates the abnormal patterning of GPe neurons during cortical SWA in dopamine-depleted mice.

**Figure 8-1.**
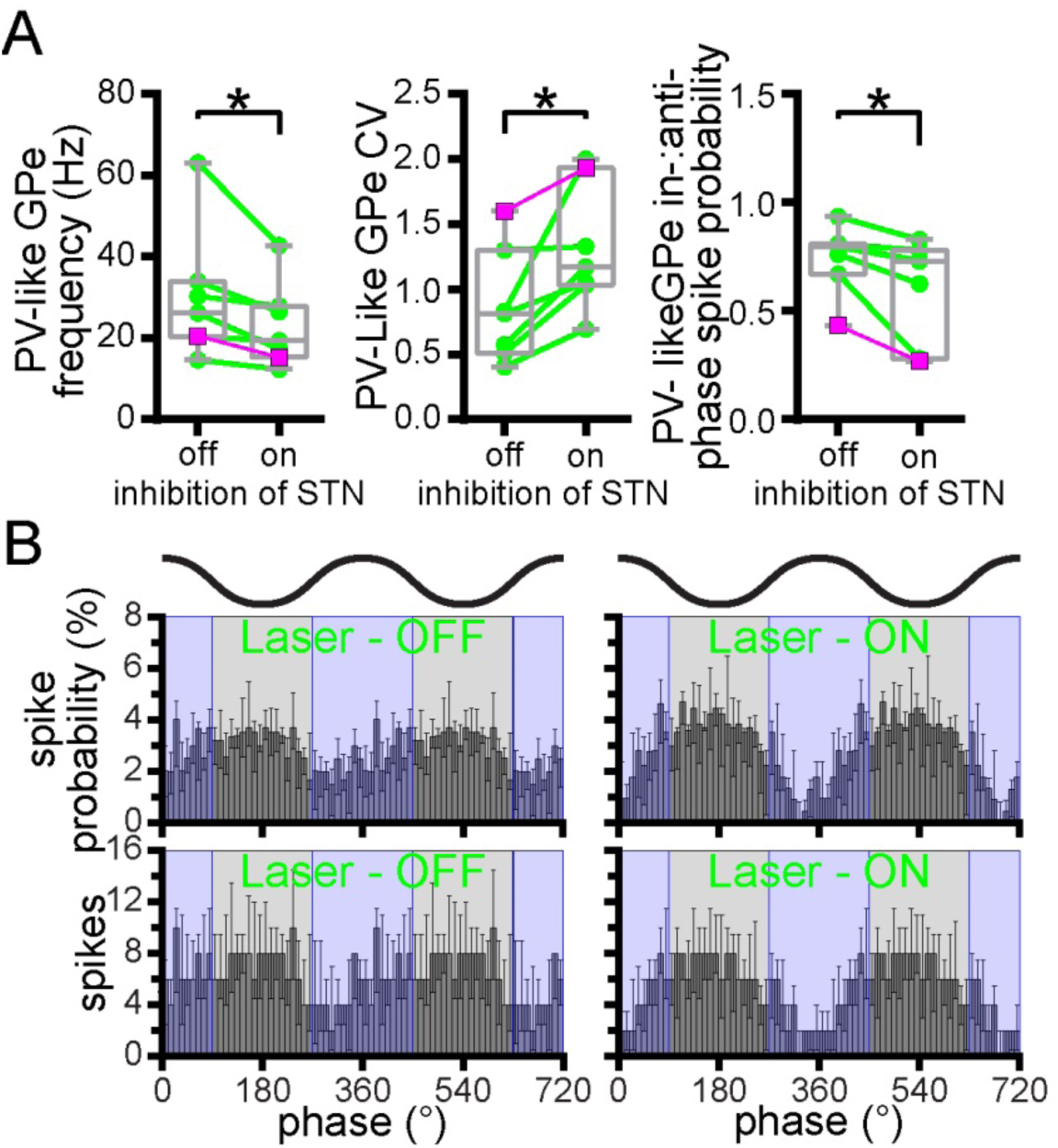
In 6-OHDA-injected mice putative PV GPe neurons were relatively uniform in their response to inhibition of STN neurons. (A-B) Optogenetic inhibition of STN neurons in 6-OHDA-injected mice 1) reduced the frequency and regularity of firing (CV) 2) reduced the in-:anti-phase spike probability of putative PV GPe neurons (A, population data, firing rate (left), CV (middle), and in-:anti-phase spike probability (right); example cell data from Fig. 8 plotted in magenta; B, population linear phase histograms). *, p < 0.05.

### Dopamine depletion reduced phase-locking of STN neuron firing to cortical SWA

In urethane-anesthetized rodents, the activity of STN neurons *in vivo* is phase-locked to the active component of SWA (Magill, Bolam, & Bevan, 2000; Magill et al., 2001; Mallet, Pogosyan, Marton, et al., 2008). To assess the impact of dopamine depletion on the rate and pattern of STN activity, 30s epochs of firing during robust SWA were compared in vehicle- and 6-OHDA-injected PV-cre mice (Table S1). Similar proportions of STN neurons were responsive to optogenetic inhibition of PV GPe neurons in vehicle and 6-OHDA treated mice (vehicle: 82 %, n = 18 of 22; 6-OHDA: 84 %, n = 21 of 25; p = 1.000; Fisher’s Exact). Dopamine depletion did not alter the firing rate (Fig. 9A-C; SWA: vehicle = 9.38, 5.90-16.4 Hz; n = 18; 6-OHDA = 11.7, 8.23-13.3 Hz; n = 21; p = 0.7122; MWU) or CV (SWA: vehicle = 1.3, 0.979-1.64; n = 18; 6-OHDA = 1.35, 1.15-1.68; n = 21; p = 0.494; MWU) of responsive STN neurons. However, there was a significant reduction in-anti-phase spike probability following dopamine depletion (Fig. 9C, D; SWA: vehicle = 3.45, 1.92-6.54; n = 18; 6-OHDA = 1.69, 0.992-3.16; n = 21; p = 0.0148; MWU). These findings are in contrast to what has been reported in dopamine-depleted rats (Magill et al., 2001; Mallet, Pogosyan, Marton, et al., 2008; Mallet, Pogosyan, Sharott, et al., 2008) and unexpected given the emergence of phase-offset prototypic GPe neuron activity, but are consistent with the downregulation of cortico-STN transmission strength in parkinsonian rodents and non-human primates (Chu et al., 2017; H. Kita & Kita, 2011a; Mathai et al., 2015; Wang et al., 2018) and downregulation in intrinsic excitability in rats and mice (McIver et al., 2019; C. L. Wilson et al., 2006; Zhu et al., 2002).

**Figure 9.**
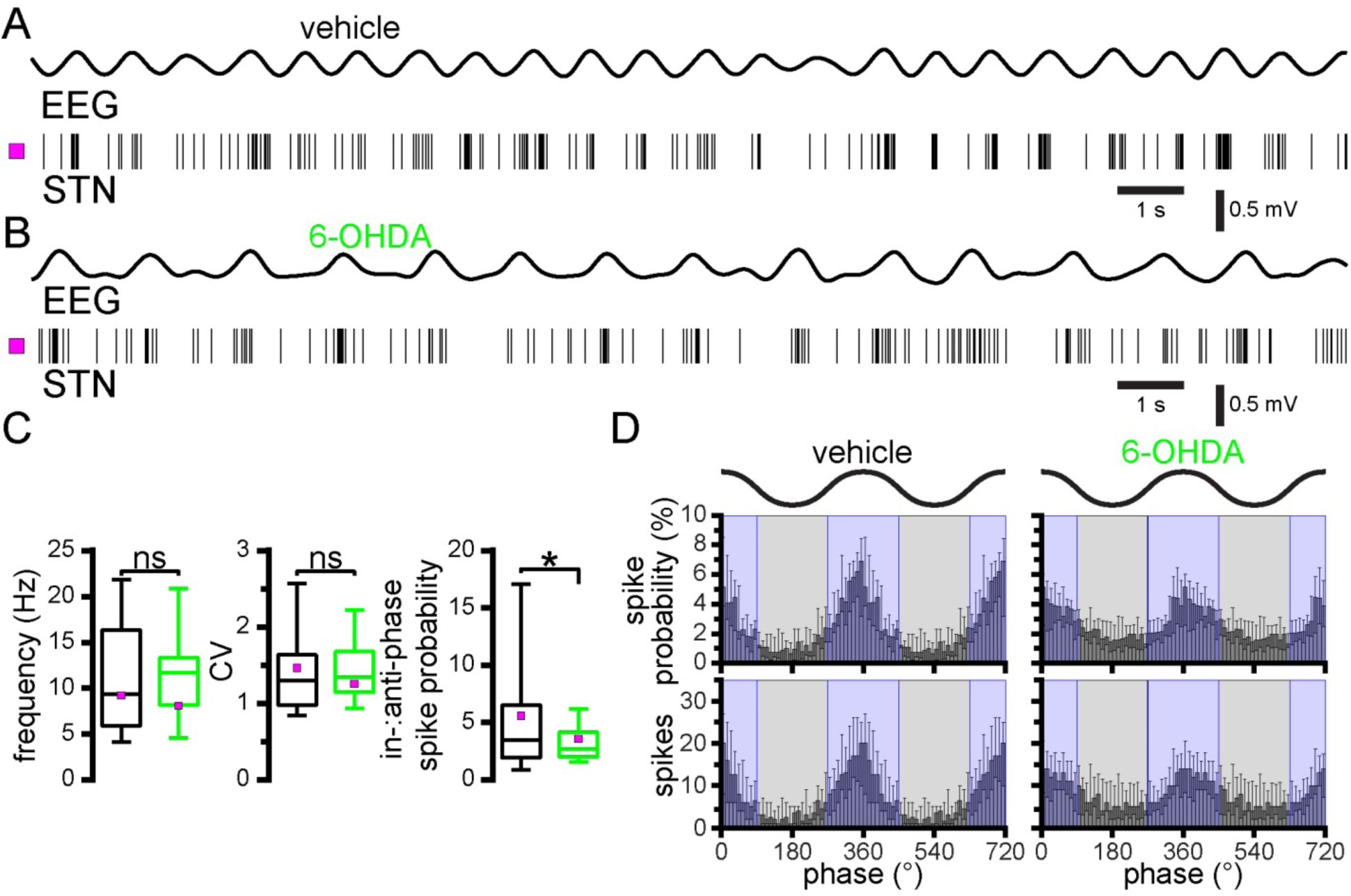
Phase-locking of STN firing to cortical SWA is reduced in 6-OHDA-injected mice. (A-D) Although STN neuron firing rate and CV were unaffected by dopamine depletion, there was a significant reduction in in-:anti-phase spike probability. (A and B) Representative examples. (C) Population data, firing rate (left), CV (middle), and in-:anti-phase spike probability (right). (D) Population linear phase histograms. *, p < 0.05. ns, not significant.

### PV GPe inhibition reduced phase-locking of STN activity to cortical SWA in control and dopamine-depleted mice

To determine how prototypic PV GPe neurons regulate STN activity in dopamine-intact and -depleted mice, the effects of optogenetically inhibiting Arch-GFP expressing PV GPe neurons for 5 seconds on STN neuron firing in control and dopamine-depleted mice were compared. The majority of responsive STN neurons were disinhibited during optogenetic inhibition of PV GPe neurons (vehicle: 72 %, n = 13 of 18; 6-OHDA: 100 %, n = 21 of 21, p = 0.6356; Fisher’s Exact). Inhibition of a minority of responsive STN neurons during PV GPe neuron silencing in vehicle-injected mice may reflect disinhibition of GPe-STN neurons due to reduced lateral inhibition in the GPe (11 %, n = 2 of 18). The failure of a subset of STN neurons to respond to PV GPe neuron inhibition presumably reflects a failure to inhibit GPe neurons presynaptic to recorded STN neurons (vehicle: 18 %, n = 4 of 22; 6-OHDA: 16 %, n = 4 of 25). Given that STN activity was not recorded at sites where PV GPe inhibition failed to elicit a response from at least one STN neuron, the proportion of STN neurons that did not respond to PV GPe inhibition is likely to be higher than that reported here. During optogenetic inhibition of PV GPe neurons, the rate and regularity of STN firing increased in both vehicle- and 6-OHDA-injected mice (Fig. 10A-C; vehicle: laser off = 11.9, 6.58-16.6 Hz; laser on = 15, 8.88-22.7 Hz; n = 18; p_h3_ = 8.424e-3; WSR; laser off CV = 1.38, 1.04-1.82; laser on CV = 0.891, 0.704-0.968; n = 17; p_h3_ = 6.27e-3; WSR; 6-OHDA: laser off =10.3, 7.30-13.65 Hz; laser on = 18.3, 12.3-29.9 Hz; n = 21; p_h4_ = 3.815e-6; WSR; laser off CV = 1.22, 0.985-1.47; laser on CV = 0.727, 0.538-0.872; n = 21; p_h4_ = 7.628-6; WSR). The firing rate of responsive STN neurons was not significantly different in dopamine-intact and -depleted mice in the absence of or during optogenetic inhibition of PV GPe neurons (Fig. 10A-C laser off: p_h1_ = 0.4990; MWU; laser on: p_h2_ = 0.4464; MWU). Although the CV of STN neuron firing was not altered by dopamine depletion alone, it was relatively reduced in 6-OHDA-injected mice during optogenetic inhibition of PV GPe neurons (Fig. 10A-C vehicle: laser off CV = 1.36, 0.936-1.79, n = 18; 6-OHDA laser off CV = 1.22, 0.985-1.47; n = 21; p_h1_ = 0.770 MWU; vehicle: laser on CV = 0.891, 0.704-0.968; n = 17; 6-OHDA: laser on CV = 0.727, 0.538-0.872; n = 21; p_h2_ = 0.02074; MWU).

**Figure 10.**
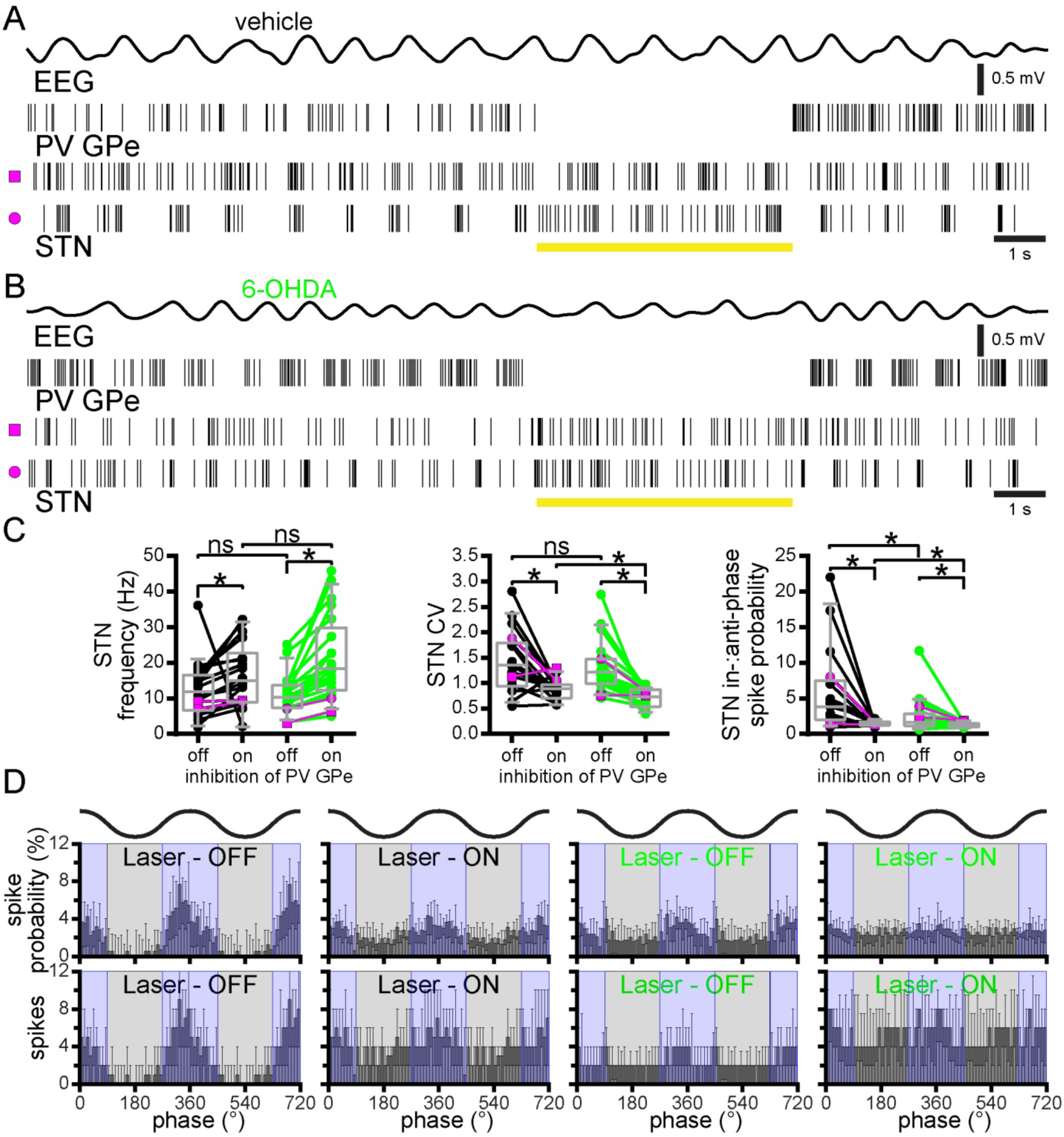
Optogenetic inhibition of PV GPe neurons disinhibits STN neurons and reduces their phase-locking to cortical SWA in both vehicle- and 6-OHDA-injected mice. (A and B) Representative examples. (C) Population data, firing rate (left), CV (middle), and in-:anti-phase spike probability (right). (D) Population linear phase histograms of STN activity relative to SWA in vehicle- (left) and 6-OHDA-injected (right) mice prior to and during optogenetic inhibition of PV GPe neurons. *p < 0.05. ns, not significant.

Inhibition of PV GPe neurons in vehicle- and 6-OHDA-injected mice also reduced the in- :anti-phase spike probability ratio (Fig. 10C, D; vehicle: laser off = 4.05, 2.02-7.75; laser on = 1.45, 1.19-1.69; n = 16; p_h4_ = 1.709e-3; WSR; 6-OHDA: laser off = 1.64, 1.13-2.80; laser on = 1.17, 0.986-1.50; n = 21; p_h3_ = 6.408e-3; WSR). In 6-OHDA injected mice, STN neurons also exhibited a reduced in-:anti-phase spike probability ratio relative to vehicle-injected controls both before and during optogenetic inhibition of PV GPe neurons (Fig. 10C, D vehicle: laser off = 3.8, 1.98-7.50; n = 17; 6-OHDA: laser off = 1.64, 1.13-2.80; n = 21; p_h2_ = 0.03024; MWU; vehicle: laser on = 1.46, 1.22-1.70; n = 17; 6-OHDA: laser on = 1.17, 0.986-1.50; n = 21; p_h1_ = 0.04414; MWU). The reduced in-:anti-phase spike probability of STN neurons in 6-OHDA mice is consistent with the downregulation of cortico-STN transmission strength in parkinsonian rodents and non-human primates (Chu et al., 2017; H. Kita & Kita, 2011a; Mathai et al., 2015; Wang et al., 2018). The additional decrease in in-:anti-phase STN activity that occurred during optogenetic inhibition of PV GPe neurons in 6-OHDA-injected mice may reflect a reduction in post-inhibitory rebound STN neuron firing (Baufreton, Atherton, Surmeier, & Bevan, 2005; Bevan, Magill, Hallworth, Bolam, & Wilson, 2002; Hallworth & Bevan, 2005) when phase-offset inhibition from the GPe was removed (Mallet et al., 2012; Mallet, Pogosyan, Sharott, et al., 2008; this study). Interestingly, optogenetic inhibition of PV GPe neurons also significantly reduced in-:anti-phase STN activity in dopamine-intact mice, arguing GPe-STN inhibition also enhances in-:anti-phase STN activity under normal conditions, despite the fact that inhibition was less anti-phasic to cortical SWA.

### Effects of tail pinch-evoked cortical ACT on D2-SPN, PV GPe neuron, and STN neuron activity in vehicle- and 6-OHDA injected mice

We next examined the activity of D2-SPNs, STN neurons, and PV GPe neurons during brief 5 second tail pinch-evoked cortical ACT (Magill et al., 2001). Previous studies in dopamine-depleted rats have utilized brief tail pinches to induce prolonged periods of cortical ACT, in which exaggerated beta band activity has been reported (Mallet et al., 2012; Mallet, Pogosyan, Sharott, et al., 2008; Mallet et al., 2019 in press). However, persistent periods of cortical activation could not be induced in mice without clear signs of inadequate anesthesia, which may explain our failure to observe exaggerated beta band activity in any of the recorded structures (Table S1; data not shown). Our study was therefore restricted to the tail pinch period. As described above, the firing rate of D2-SPNs during SWA was elevated in 6-OHDA-versus vehicle-injected mice (Fig. 11A-C; vehicle: SWA = 2.5, 1.60-3.40 Hz; n = 34; 6-OHDA: SWA = 3.4, 2.10-7.80; n = 21; p_h2_ = 0.02476; MWU). In response to ACT, the firing rate of D2-SPNs in vehicle-injected mice decreased (Fig. 11A-C; vehicle: ACT = 0.300, 0.00-1.35 Hz; n = 34; p_h3_ = 1.751e-3; WSR), in contrast to D2-SPNs in 6-OHDA-injected animals, which maintained their elevated firing rate (Fig. 11; 6-OHDA: ACT = 4.00, 2.10-9.60 Hz; n = 21; p_h1_ = 0.5563; WSR). As a result the firing of D2-SPNs during tail pinch-evoked cortical ACT was greater in 6-OHDA mice (MWU; p_h4_ = 2.656e-4). These data are consistent with a recent study in rats (Sharott et al., 2017) in which the firing of D2-SPNs was also elevated in dopamine-depleted rats during more prolonged periods of ACT.

**Figure 11.**
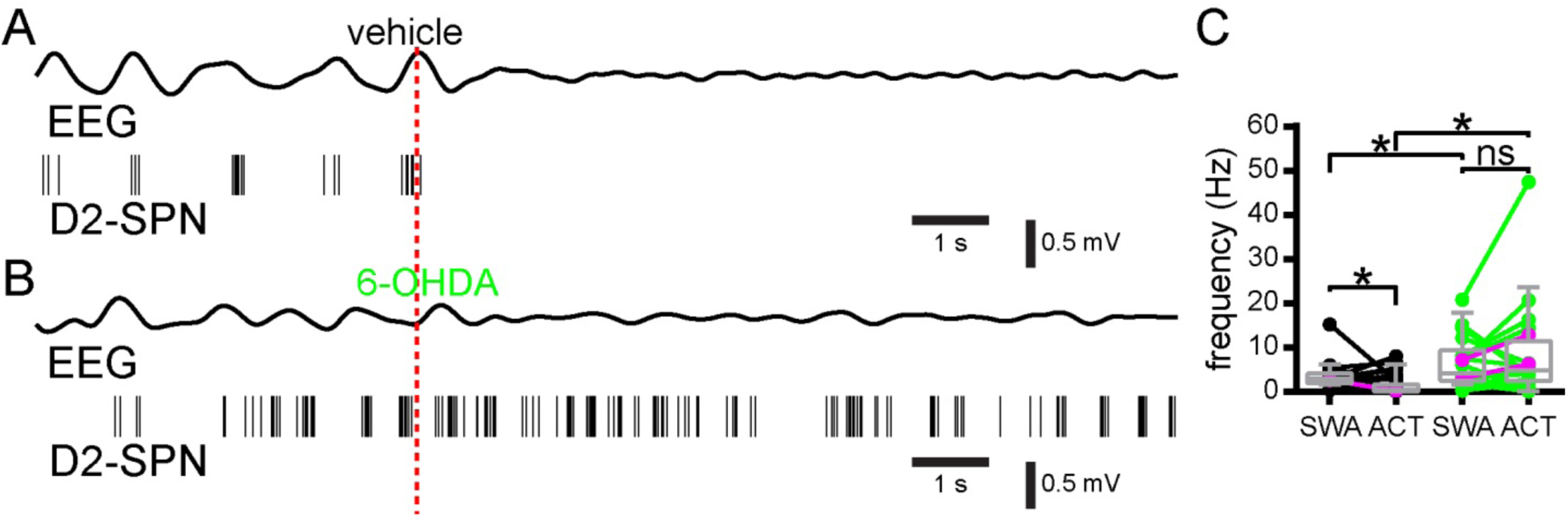
Tail pinch-evoked cortical ACT suppresses D2-SPN activity in vehicle-but not 6-OHDA injected mice. (A-C) In vehicle-injected, dopamine-intact mice D2-SPN activity decreased during tail pinch-evoked cortical activation (ACT) relative to SWA. In 6-OHDA-injected, dopamine-depleted mice D2-SPN activity was similar during SWA and ACT. Thus the firing rate of D2-SPNs during cortical ACT was greater in 6-OHDA-injected mice. (A, B) Representative examples (onset of tail pinch denoted by dashed red line). (C) Population data, examples plotted in magenta. *, p < 0.05. ns, not significant.

Dopamine depletion altered the response of optogenetically identified PV GPe neurons to ACT in a manner that was consistent with earlier studies of prototypic neurons in rats (Fig. 12A-C) (Abdi et al., 2015; Magill et al., 2001; Mallet et al., 2012; Mallet, Pogosyan, Marton, et al., 2008). In vehicle-injected control mice, the firing of PV GPe neurons significantly increased during ACT (Fig. 12A, C; vehicle: SWA = 15.8, 9.00-42.0 Hz; ACT = 27.2, 18.6-41.0 Hz; n = 23; p_h2_ = 0.02512; WSR). In contrast, the firing of PV GPe neurons decreased during ACT in parkinsonian mice (Fig. 12B, C; 6-OHDA: SWA = 15.6, 12.4-19.0 Hz; ACT = 1.8, 0.20-16.0 Hz; n = 15; p_h3_ = 3.48e-3; WSR). As a result, the frequency of PV GPe neuron firing was dramatically reduced in dopamine-depleted mice relative to control mice during ACT (Fig. 12C ACT: p_h4_ = 3.27e-5; MWU). However, in contrast to previous studies in rats (Magill et al., 2001; Mallet et al., 2012; Mallet, Pogosyan, Marton, et al., 2008) STN neurons in both vehicle- and 6-OHDA-injected mice exhibited no difference in firing rate during cortical SWA and a similar increase in firing during ACT (Fig. 12 D-F; vehicle: SWA = 13.8, 10.0-17.8 Hz; ACT = 18.8, 12.2-23.0; n = 15; p_h3_ = 0.03552; WSR; 6-OHDA: SWA = 9.60, 6.80-13.6 Hz; ACT = 15.6, 9.40-18.4 Hz; n = 15; p_h4_ = 0.01221; WSR) (SWA: p_h2_ = 0.1743; MWU; ACT: p_h1_ = 0.2715; MWU). As for cortical SWA, these data are consistent with increased striatopallidal transmission in dopamine-depleted mice, which is not fully reflected in the activity of STN neurons, possibly due to adaptive changes in that nucleus.

**Figure 12.**
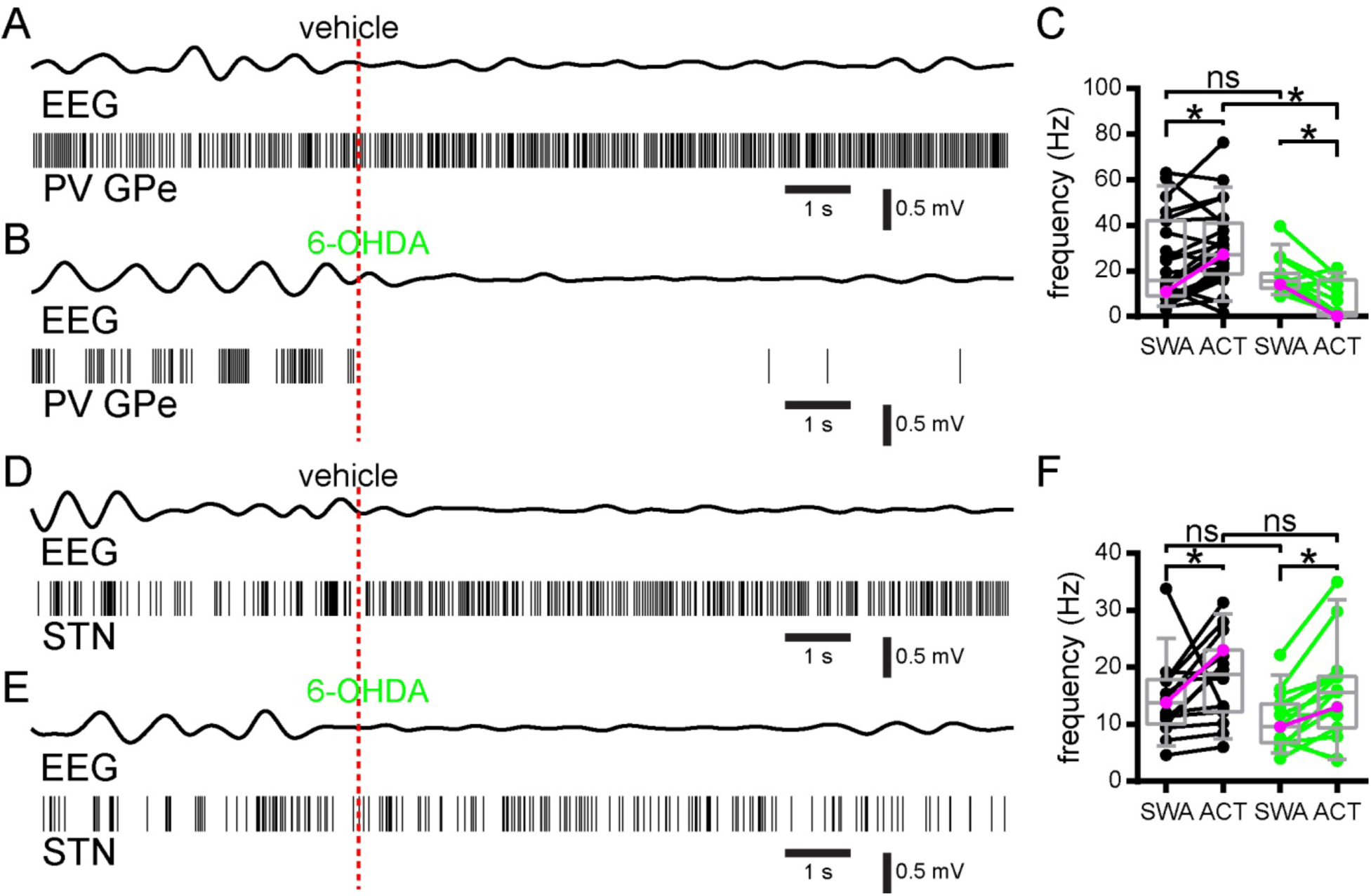
Effect of tail pinch-evoked cortical ACT on PV GPe and STN neuronal activity in control and dopamine depleted mice. (A-C) In vehicle-injected dopamine-intact mice cortical ACT increased PV GPe neuron activity relative to SWA. In 6-OHDA-injected, dopamine-depleted mice PV GPe neuron activity was suppressed during cortical ACT. (A, B) Representative examples (onset of tail pinch denoted by dashed red line). (C) Population data. (D-F) STN neuron activity significantly increased during cortical ACT relative to SWA in both vehicle- and 6-OHDA-injected animals. However, the rate of STN activity during ACT was similar in dopamine-intact and -depleted mice. (D, E) Representative examples (onset of tail pinch denoted by dashed red line). (F) Population data, examples plotted in magenta. *, p < 0.05. ns, not significant.

## Discussion

Although brain circuit activity under anesthesia does not recapitulate activity in awake behaving animals, this preparation enabled us to study 1) how dopamine depletion alters basal ganglia activity during multi-second periods of stereotyped synchronous cortical activity, which is informative given the emergence of excessively correlated cortico-basal ganglia-thalamocortical activity in PD 2) basal ganglia activity in the absence of movement-associated proprioceptive feedback, which given the perturbation of movement kinematics in Parkinsonism helps to restrict the potential sources of abnormal firing. By combining this recording configuration with optogenetic identification and inhibition, we were able to not only measure the activity of specific classes of neuron in the basal ganglia but also to determine how each neuron type influenced its targets. We demonstrated in 6-OHDA-injected mice that hyperactive D2-SPNs were responsible for prototypic PV GPe neuron activity that was relatively anti-phasic to cortical SWA. Downregulation of autonomous prototypic PV GPe neuron firing was not observed following the loss of dopamine, and therefore cannot account for the emergence of pauses in PV GPe neuron activity *in vivo*. Despite the development of more temporally offset GPe and STN activity in 6-OHDA-injected mice, the STN did not exhibit elevated firing during SWA. STN output also opposed rather than facilitated abnormal GPe activity. Finally, during sustained cortical ACT in 6-OHDA-injected mice, D2-SPNs exhibited hyperactivity and PV GPe neurons exhibited hypoactivity, but the rate of STN activity was unchanged. Together, these data argue that in dopamine-depleted mice increased striatopallidal inhibition is largely responsible for abnormal prototypic PV GPe neuron activity and that despite being disinhibited, the STN does not exhibit hyperactivity, presumably due to recently reported cellular and synaptic adaptations.

Our finding that D2-SPNs discharge excessively during cortical SWA or ACT in dopamine-depleted mice relative to controls is consistent with 1) the loss of inhibitory dopaminergic modulation of cortico-D2-SPN transmission and D2-SPN excitability (Gerfen & Surmeier, 2011) 2) D2-SPN hyperactivity in the majority of studies in anesthetized rats (Ballion et al., 2009; Escande et al., 2016; Mallet et al., 2006; Parker et al., 2018; Ryan, Bair-Marshall, & Nelson, 2018; Sharott et al., 2017), and in awake, immobile mice (Parker et al., 2018; Ryan et al., 2018). Excessive firing of D2-SPNs in immobile dopamine-depleted mice but not during movement could indicate that movement-associated D2-SPN activity is limited by reduced cortical drive and/or abnormal proprioceptive feedback, despite the best attempts of experimenters to control for these issues. Another possibility is that adaptive alterations in cortico-D2-SPN synaptic transmission and connectivity and the intrinsic excitability of D2-SPNs underlie the absence of movement-related D2-SPN hyperactivity and ensemble activity in dopamine-depleted mice (Day et al., 2006; Escande et al., 2016; Fieblinger et al., 2014; Mallet et al., 2006; Taverna et al., 2008). However, why homeostatic plasticity would not also limit cortical driving of D2-SPNs during cortical SWA or ACT, or during periods of awake immobility remains unclear (Parker et al., 2018; Ryan et al., 2018).

Under normal conditions, GPe-STN network activity is largely irregular and de-correlated (Arkadir, Morris, Vaadia, & Bergman, 2004; Elias et al., 2007; Magill et al., 2000; Matsumura, Kojima, Gardiner, & Hikosaka, 1992; Wichmann, Bergman, & DeLong, 1994). In PD and its models GPe and STN neurons exhibit increases in pauses in firing and/or burst discharge, the occurrence of which may be correlated and coherent with cortical activity (Delaville et al., 2015; Magill et al., 2001; Mallet, Pogosyan, Sharott, et al., 2008; Quiroga-Varela et al., 2013; Sanders et al., 2013; Tachibana et al., 2011; Walters et al., 2007). To determine how abnormal D2-SPN activity affects the GPe-STN network, we first examined the effects of dopamine depletion on PV GPe neurons, a subset of prototypic GPe neurons that represents over half of all GPe neurons, approximately 85% of prototypic GPe neurons, and the vast majority of STN projecting neurons (Abdi et al., 2015; Dodson et al., 2015; Hernandez et al., 2015). We found that following dopamine depletion, PV GPe neuron firing was relatively anti-phasic to cortical SWA, as previously demonstrated in rats (Abdi et al., 2015; Mallet, Pogosyan, Sharott, et al., 2008; Walters et al., 2007; Zold, Larramendy, Riquelme, & Murer, 2007), but there was no change in firing rate, consistent with the majority of these reports (Abdi et al., 2015; Walters et al., 2007; Zold et al., 2007).

Given the phase offset of GPe neuron activity in dopamine-depleted rodents relative to cortical SWA, D2-SPN, and STN activity, hyperactive D2-SPNs were proposed to be responsible for increased anti-phasic GPe neuron activity (Nevado-Holgado, Mallet, Magill, & Bogacz, 2014; Walters et al., 2007). Consistent with this idea, pharmacological manipulations, such as intrastriatal delivery of the NMDAR antagonist D-APV (Zold et al., 2012) or the GABA_A_R agonist muscimol (H. Kita & Kita, 2011a), to suppress striatal activity and output, diminished anti-phasic GPe activity. Given that D1-SPNs also provide a substantial collateral projection to the GPe (Cazorla et al., 2014; Fujiyama et al., 2011; Kawaguchi, Wilson, & Emson, 1990; Wu, Richard, & Parent, 2000), it is possible that they too contribute to abnormal anti-phasic GPe activity. However, D1-SPN activity is suppressed in dopamine-depleted mice, arguing that their inhibition of GPe neurons is likely to be less significant than that mediated by D2-SPNs (Escande et al., 2016; Mallet et al., 2006; Parker et al., 2018). In this study, we demonstrate that selective optogenetic inhibition of D2-SPNs alone is sufficient to rapidly and reversibly reduce anti-phasic GPe neuron activity in dopamine-depleted mice, in a manner that was precisely time-locked to the 5 second period of optogenetic manipulation. Thus, given their elevated firing and transmission probability, D2-SPNs are likely to be the primary generators of anti-phasic GPe activity in the dopamine-depleted state (Escande et al., 2016; Lemos et al., 2016; Miguelez et al., 2012; Parker et al., 2018; Ryan et al., 2018; Zold et al., 2012). That said, anti-phasic GPe activity was not completely eliminated, presumably because the presynaptic D2-SPN population was only partially inhibited due to the spatially restricted nature of optogenetic stimulation and <100 % transfection of D2-SPNs with Arch-GFP. PV GPe neurons also exhibited dramatic hypoactivity during sustained cortical ACT in dopamine-depleted mice, which is consistent with the relatively elevated and persistent activity of D2-SPNs in this brain state, as reported here and in analogous studies of identified striatopallidal neurons (Sharott et al., 2017) and unidentified (Magill et al., 2001; Mallet et al., 2008) and identified (Abdi et al., 2015) PV GPe neurons in rats.

Previous studies have suggested that following the loss of dopamine, downregulation of GPe HCN channels causes a reduction in their autonomous firing, which leaves GPe neurons susceptible to entrainment by synaptic inputs (Chan et al., 2011; Shouno et al., 2017). However, another study reported that only Npas1 GPe neurons, the majority of which are arkypallidal neurons that project to the striatum, exhibit downregulated activity (Hernandez et al., 2015). Here, we found that the autonomous activity of prototypic PV GPe neurons was significantly upregulated in 6-OHDA-injected mice, consistent with upregulated autonomous and post-inhibitory rebound firing in a subset of unidentified GPe neurons in 6-OHDA-injected rats (Miguelez et al., 2012). The loss of intrinsic activity cannot therefore be responsible for the relatively interrupted activity of PV GPe neurons *in vivo* in dopamine-depleted mice. The upregulation of autonomous firing in PV GPe neurons presumably represents a homeostatic response to elevated D2-SPN-GPe transmission in dopamine-depleted mice, which could contribute to the maintenance of PV GPe neuron firing rates during cortical SWA. That said, upregulated autonomous firing was clearly insufficient to prevent PV GPe neuron hypoactivity during sustained cortical ACT and D2-SPN hyperactivity. The mechanisms underlying the upregulation of autonomous PV GPe neuron firing have yet to be elucidated but could be triggered by alterations in the activation of postsynaptic Ca_v_1 channels that report spiking activity (Chan et al., 2011) or NMDARs that report the strength of synaptic excitation (McIver et al., 2019).

In urethane-anesthetized rodents STN neurons discharge in phase with cortical SWA due to direct cortical excitation (Magill et al., 2000, 2001; Mallet, Pogosyan, Sharott, et al., 2008). Despite the emergence of more anti-phasic activity in prototypic PV GPe neurons in dopamine-depleted mice and disinhibition of STN neurons at the time that they are receiving cortical excitation, the activity of STN neurons in phase with cortical SWA was not elevated. The mean frequency of STN activity in vehicle- and 6-OHDA-injected mice was similar and the ratio of in- to anti-phase STN activity decreased modestly but significantly in dopamine-depleted mice. A similar picture emerged during cortical ACT, although the frequency of PV GPe neuron activity was dramatically reduced in 6-OHDA-injected mice, the frequency of STN activity was similar in dopamine-intact and -depleted mice. These data are in contrast to some (Deffains et al., 2016; Delaville et al., 2015; Magill et al., 2001; Mallet, Pogosyan, Sharott, et al., 2008; Soares et al., 2004; Tachibana et al., 2011; Walters et al., 2007; Wichmann, Kliem, & Soares, 2002) but not other (Ni et al., 2001; Ryu et al., 2011) *in vivo* recording studies of STN activity in PD models. Why is the STN not hyperactive in Parkinsonian mice when the frequency of presynaptic PV GPe neuron firing is reduced (Atherton, Menard, Urbain, & Bevan, 2013)? One possible explanation is the engagement of homeostatic synaptic and cellular plasticity triggered by the loss of dopamine. Thus, following degeneration of midbrain dopamine neurons, GPe-STN inputs proliferate (Chu et al., 2015; Fan et al., 2012) and cortico-STN inputs are lost (Chu et al., 2017; Mathai et al., 2015; Wang et al., 2018), leading to an approximately 4-fold reduction in the strength of synaptic excitation versus inhibition in the STN. Furthermore, the autonomous activity of STN neurons is downregulated (McIver et al., 2019). These alterations are NMDAR-dependent and can be generated in dopamine-intact mice through chemogenetic activation of D2-SPNs and subsequent disinhibition of the STN or through application of exogenous NMDA *ex vivo* (Chu et al., 2015; Chu et al., 2017; McIver et al., 2019). Furthermore, in dopamine-depleted mice, plasticity induced by chemogenetic activation of D2-SPNs is occluded (Chu et al., 2017; McIver et al., 2019). Thus, following the loss of dopamine, hyperactivity of D2-SPNs leads to reduced firing of GPe neurons at the time the STN receives synaptic excitation from hyperdirect pathway cortical inputs, which leads to increased activation of STN NMDARs that in turn triggers compensatory synaptic and cellular adaptations. As a result, the impact of cortical excitation on STN activity is downregulated both *ex vivo* and *in vivo* (Chu et al., 2017; H. Kita & Kita, 2011a; Mathai et al., 2015; this study). It has been suggested that enhanced hyperdirect pathway patterning contributes to oscillatory entrainment of the basal ganglia in PD and its models (Moran et al., 2011; Pavlides, Hogan, & Bogacz, 2015; Tachibana et al., 2011) but our results and recent studies that demonstrate that cortico-STN input is strongly downregulated in mice, rats, and monkeys (Chu et al., 2017; H. Kita & Kita, 2011a; Mathai et al., 2015; Wang et al., 2018), argue against this thesis. Indeed, if hyperdirect cortical input was sufficient to pattern Parkinsonian STN activity it should be more apparent during inhibition of prototypic PV GPe neurons, but in fact we found the opposite to be the case. In 6-OHDA-injected mice, inhibition of PV GPe neurons almost entirely eliminated phase locking of STN activity to cortical SWA, consistent with the idea that patterning of STN activity by the indirect pathway relative to the hyperdirect pathway is elevated in Parkinsonism.

The reciprocally connected GPe-STN network has been suggested to be a contributor to pathologically correlated cortico-basal ganglia-thalamocortical activity in PD (Moran et al., 2011; Shouno et al., 2017; Tachibana et al., 2011). Indeed, lesion (Bergman, Wichmann, & DeLong, 1990), electrical stimulation (Benabid, Chabardes, Mitrofanis, & Pollak, 2009; Wichmann, Bergman, & DeLong, 2018), pharmacological (Levy et al., 2001; Tachibana et al., 2011), optogenetic (Mastro et al., 2017; Yoon et al., 2014; Mallet et al., 2019 in press), genetic (Chu et al., 2017; Zhuang et al., 2018), and chemogenetic (Assaf & Schiller, 2019; McIver et al., 2019) manipulation of the GPe and/or STN have demonstrated that altering their pathological activity can ameliorate motor dysfunction in PD and its models. Interestingly, we found that optogenetic inhibition of STN activity facilitated rather than ameliorated abnormal anti-phasic GPe neuron activity during SWA, arguing that STN-GPe transmission *opposed* the abnormal patterning of the GPe by indirect pathway D2-SPNs in dopamine-depleted mice. Although the rate and pattern of STN activity were not significantly altered by dopamine depletion, the emergence of temporally offset prototypic GPe activity may cause the GPe-STN network to more powerfully synchronize basal ganglia output neurons in PD (Lobb & Jaeger, 2015; Mastro et al., 2017; Willard et al., 2019). Thus, increased patterning of GPi/SNr could occur through alternating bouts of concurrent 1) GPe inactivity and STN activity, both of which should elevate basal ganglia output; 2) GPe activity and STN inactivity, which should inhibit basal ganglia output. Indeed, manipulations that oppose the patterning of GPe (and STN) neurons by D2-SPNs e.g. optogenetic activation of PV GPe neurons (Mastro et al., 2017), chemogenetic activation of the GPe (Assaf & Schiller, 2019), and chemogenetic and genetic manipulation of the STN (Chu et al., 2017; Zhuang et al., 2018; McIver et al., 2019) ameliorate motor dysfunction in PD models.

## Materials and Methods

### Statistical analysis

Statistics were performed using Prism 6 (GraphPad Software, Inc., La Jolla, CA, USA) or R (https://www.r-project.org/; exactRankTests package). Population data are expressed as median and interquartile range and illustrated using box plots (central line: median; box: 25 % - 75 %; whiskers: 10 % - 90 %); outliers were not excluded from analysis. Paired data are illustrated using line plots. It should be noted that some data are excluded from paired analyses that are included in unpaired analyses (e.g. when CV could not be calculated due to an absence of firing); these data are represented in box plots overlying paired line plots. Because no assumptions were made concerning the distribution of population data, non-parametric statistics were used throughout. Paired and unpaired data were compared using the non-parametric Mann-Whitney U (MWU) and Wilcoxon Signed Rank (WSR) tests, respectively, and Fisher’s exact test was used for contingency analyses. When applicable, the Holm-Bonferroni correction for multiple comparisons (Holm, 1997) was applied (adjusted p-values are indicated as p_h#_, where # is the adjustment factor). P values < 0.05 were considered significant. For the primary findings reported in the manuscript, sample sizes for Mann-Whitney and Wilcoxon tests were estimated to achieve a minimum of 80% power using formulae described by (Noether, 1987). The effect sizes used in these power calculations were estimated using data randomly drawn from uniform distributions (runif() function in R stats package). For Mann-Whitney tests, with a 50 percentile change in median between groups X and Y (the interquartile ranges of the groups don’t overlap) P(Y > X) ≈ 0.88 giving an estimation that at least 10 observations per group would be needed to achieve 80% power; for a 25 percentile change (the median of Y falls outside the interquartile range of X) P(Y > X) ≈ 0.72 and the estimated requirement is at least 27 observations per group. For Wilcoxon tests, if all pairs of observations show the same direction of change, P(X + X’ > 0)= 1 giving an estimation that at least 10 observations would be needed to achieve 80% power (note though that it is possible to show empirically that 6 observations gives 100% power in this case); if 90% of observations show the same direction of change, P(X + X’ > 0) ≈ 0.98 and the estimated requirement is at least 12 pairs of observations

### Animals

Experiments involving mice were carried out in compliance with the Northwestern University and University of Bordeaux Institutional Animal Care and Use Committees and NIH and European Economic Community guidelines. Adult male mice including A2A-cre (Tg(Adora2a-cre)KG139Gsat; 163, 104-196 days old; n = 18), PV-cre (B6.Cg-Pvalb^tm1.1(cre)Aibs^/J; 151, 126-167 days old; n = 14), PV-cre X Ai9 mice (B6.129P2-Pvalb^tm1(cre)Arbr^/J X B6.Cg-Gt(ROSA)26Sor^tm9(CAG-tdTomato)Hze^/J; 69.5, 67-71 days old; n = 14), and GABRR3-cre (Tg(Gabrr3-cre)KC112Gsat; 169, 68-293 days old; n = 7) mice were used. Mice were housed under a 14h/10h (*in vivo* experiments) or 12h/12h (*ex vivo* experiments) light/dark cycle with food and water *ad libitum*, received regular veterinary inspections, and underwent only those procedures detailed here and tail-clipping for the purpose of genotyping. Experiments were performed during the light cycle.

### Unilateral 6-OHDA/vehicle and AAV injection

6-OHDA/vehicle and AAVs were injected using a stereotaxic instrument (Neurostar, Tubingen, Germany; David Kopf Instruments, Tujunga, CA, USA). Anesthesia was induced with vaporized 3-4 % isoflurane (Smiths Medical ASD, Inc., Dublin, OH, USA; Tem-Sega, Inc., Pessac, France) before injection of ketamine/xylazine (75/10 or 87/13 mgkg^-1^, IP). Within 2-5 minutes, desipramine (25 mg/kg, IP) and pargyline (50 mg/kg, IP) were injected to enhance the selectivity and toxicity of intracerebral 6-OHDA injections, respectively. After securing the animal in the stereotaxic instrument, anesthesia was maintained with 1-2 % isoflurane. 1-2 µl of 6-OHDA (3-5 mg/ml) in HEPES buffered saline (HBS (in mM): 140 NaCl, 23 glucose, 15 HEPES, 3 KCl, 1.5 MgCl_2_, 1.6 CaCl_2_; pH 7.2 with NaOH; 300–310 mOsm/L) plus 0.02 % ascorbate or vehicle (HBS plus 0.02 % ascorbate) was then injected into the MFB (from Bregma: AP, −0.7 mm; ML, 1.2 mm; DV, 4.7 mm) over a 10 minute period. The injectate was then allowed to diffuse for a further 10 minutes prior to retraction of the syringe. For D2-SPN, GPe, or STN neuron identification/silencing experiments in which A2A-cre, PV-cre, or GABRR3-cre transgenic mice were used, AAV expressing AAV9.CBA.Flex.Arch-GFP.WPRE.SV40 (Addgene viral prep # 22222-AAV9; Chow et al., 2010) was injected into each target structure (striatum: 4 x 300 nl; AP, 0.4 mm and 0.9 mm; ML, 2.2 mm; DV, 3.7 mm and 2.7 mm; GPe: 2 x 300 nl; AP, −0.27 mm; ML, 2 mm; DV, 3.95 mm and 3.45 mm; STN: 500 nl; AP, −2.0 mm; ML, 1.6 mm; DV, 4.65 mm). Each AAV injection took place over 5 minutes, followed by a further 5 minute period prior to syringe retraction.

### In vivo electrophysiological recording

2-4 weeks following surgery, mice were anesthetized with 3-4 % isoflurane, injected with urethane (1.25 g/kg, IP), allowed to rest in their homecage for 1 hour, and then injected with ketamine/xylazine (16/0.8 mg/kg, IP) every 10-20 minutes until the toe-pinch withdrawal reflex was abolished. Mice were placed into a stereotaxic instrument (David Kopf Instruments) for the duration of the recording session with ketamine/xylazine supplements administered as needed to maintain anesthesia. Craniotomies were drilled over the ipsilateral motor cortex (AP, 1.4 mm; ML, 1.5 mm) and two recording sites among the striatum (AP, 0.5 mm; ML, 2.5 mm), GPe (AP, −0.3 mm; ML, 2.0 mm), and STN (AP, −1.7 mm; ML, 1.6 mm) depending on the experiment. A peridural screw “electrode” (MS-51960-1; McMaster-Carr, Chicago, IL, USA) was implanted over primary motor cortex from which the intracranial electroencephalogram (EEG) was recorded. Extracellular single unit recordings were acquired using silicon tetrode arrays (A1x4-tet-10mm-100-121-A16; NeuroNexus Technologies, Ann Arbor, MI, USA) connected to a 64-channel Digital Lynx (Neuralynx, Boseman, MT, USA) data acquisition system with a unity gain headstage, at a sampling frequency of 40 kHz, a gain of 14 X, with reference wire implanted opposed to the ipsilateral temporal musculature. Online digital FIR filters were applied. Single unit activity was band pass filtered between 200-9000 Hz and LFP and EEG signals were band pass filtered between 0.1-400 Hz. Optogenetic stimulation was delivered using a custom 577 nm laser system (Genesis MX STM 577-500 OPSL CW; Coherent Inc., Santa Clara, CA, USA) that was fiber coupled to an optrode with an identical array of tetrodes (A1x4-tet-10mm-100-121-OA16; NeuroNexus Technologies). Silicon probes were dipped in a lipophilic florescent dye (DiI; 20 mg/ml in 50 % acetone/methanol; D282; ThermoFisher Scientific, Waltham, MA, USA) prior to initial penetration to identify recording sites in histological sections (Fig. S1).

Unit activity, LFP, and EEG were simultaneously recorded for several minutes during cortical slow wave activity (SWA). This was followed by at least two periods of optogenetic stimulation of Arch (< 6 mW) for 5 seconds that were separated by at least three minutes. Laser intensity was calibrated as resultant power from the optrode fiber tip prior to probe implantation and verified at the conclusion of each experiment. Activated cortical states (ACT) were evoked by tail pinch using a custom pneumatic device. After SWA had stabilized following ACT, another tail pinch was applied. This sequence was repeated until at least three trials had been recorded or units were lost.

### Histological processing of in vivo tissue

Following recording, mice were given a lethal dose of anesthetic and transcardially perfused with saline for 2 minutes followed by 4 % paraformaldehyde in 0.1 M PB, pH 7.4 for approximately 20 minutes. The brain was then removed, held in the same fixative overnight, and then washed in 0.01 M phosphate-buffered saline, pH 7.4 (PBS; P3813; Millipore Sigma, Darmstadt, Germany) before being sectioned in the coronal plane at 70 µm using a vibratome (VT1000S; Leica Microsystems Inc., Richmond, Illinois, USA). Sections were then washed 3 times in PBS before incubation for 48-72 hr at 4 °C in a mixture of PBS, 0.5 % Triton X-100 (T8787; Millipore Sigma), and 2 % normal donkey serum (017-000-121; Jackson ImmunoResearch, West Grove, PA, USA; PBS-T) containing primary antibodies (see below). Sections were then washed in PBS before incubation for 90 minutes at room temperature (RT) in PBS-T containing secondary antibodies (see below). Lastly, sections were washed in PBS and mounted on slides with Prolong Gold Antifade Reagent (P36930; ThermoFisher Scientific, Waltham, MA, USA). Mountant was allowed to cure for at least 24 hours prior to storage at 4 °C or imaging. GFP, DiI, and immunofluorescent labelling were imaged using a Zeiss Axioskop 2 microscope (Zeiss, Oberkochen, Germany), an AxioCam CCD camera (426508-9901-000; Zeiss), and Neurolucida software (MFB Bioscience, Williston, VT, USA). Representative images were acquired using confocal laser scanning microscopy (A1R; Nikon, Melville, USA).

A 1:6 series of the striatum and substantia nigra was processed for the immunohistochemical detection of tyrosine hydroxylase (TH; 1:500 mouse anti-TH; MAB318, Millipore Sigma; 1:250 Alexa Fluor 488 donkey anti-mouse IgG; 715-545-152; Jackson Immunoresearch) and expression was quantified, as described previously (Fan et al., 2012). Immunoreactivity was averaged across three evenly spaced rostral, middle, and caudal sections. Cortical immunoreactivity was subtracted from striatal immunoreactivity to normalize for background fluorescence. Dopamine depletion was assessed from the immunoreactivity in the vehicle or 6-OHDA injected hemisphere expressed as a percentage of immunoreactivity in the contralateral hemisphere (vehicle: ipsilateral striatal TH immunoreactivity = 102, 97-104 %; n = 16; 6-OHDA: ipsilateral striatal TH immunoreactivity = −3, −8-0 %; n = 23; p = 5.303e-11; MWU). For PV-cre mice, in which PV GPe neurons expressed Arch-GFP, adjacent sections of the GPe were processed for the immunohistochemical detection of PV (1:200 rabbit anti-PV; 195002; or 1:200 guinea pig anti-PV; 195004; Synaptic Systems, Goettingen, Germany; 1:250 Alexa Fluor 594 donkey anti-rabbit IgG; 711-585-152; or 1:250 Alexa Fluor 594 donkey anti-guinea pig IgG; 706-585-148; Jackson Immunoresearch) or FoxP2 (1:500 rabbit anti-FoxP2; HPA000383; Millipore Sigma; 1:250 Alexa Fluor 594 donkey anti-rabbit IgG; 711-585-152; Jackson Immunoresearch). Sections of the STN from PV-cre and GABRR3-cre mice, the GPe from GABRR3-cre and A2A-cre mice, and the striatum from A2A-cre mice were processed for the immunohistochemical detection of the neuronal markers NeuN (1:200 mouse anti-NeuN; MAB377; Millipore Sigma; or 1:1,000 rabbit anti-NeuN; EPR12763; Abcam, Cambridge, UK; 1:250 Alexa Fluor 594 donkey anti-mouse IgG; 715-585-150; or 1:250 Alexa Fluor 594 donkey anti-rabbit IgG; 711-585-152; Jackson Immunoresearch) or HuC/D (1:66 mouse anti-HuC/D; A-21271; ThermoFisher Scientific; 1:250 Alexa Fluor 594 donkey anti-mouse; 715-585-150; Jackson Immunoresearch) to aid reconstruction of electrode tracks and sites of recording (Fig. S1).

Stereological counts of Arch-GFP, PV, and FoxP2 expressing GPe cells were determined using the optical dissector method (West, Slomianka, & Gundersen, 1991) on a 1:4 series of sections. Structures were traced with a 10 X objective (420943-900-000; Zeiss, Oberkochen, Germany) and imaged using a 40 X oil- (000000-1022-818; Zeiss, Oberkochen, Germany) or 60 X water-immersion (UplanApo 60X/1.2 NA; Olympus, Tokyo, Japan) objectives. Counting frames of 100 x 100 or 90 x 90 µm, and grid sizes of 300 x 300 or 200 x 200 µm, respectively were used. Images were taken at 1 µm intervals for 5 µm beneath a 1 µm guard zone (Neurolucida system, MFB Bioscience, Williston, VT, USA).

### In Vivo Electrophysiological Analysis

Putative single unit activity was discriminated with Plexon Offline Sorter software (Version 3; Plexon. Inc., Dallas, TX) using a combination of template matching, principal component analysis, and manual clustering. Spike times were aligned to the peak of the extracellularly recorded action potential and a dead time of 500 µs was utilized during discrimination of units (Adamos, Laskaris, Kosmidis, & Theophilidis, 2010; Lu et al., 2016; Maccione et al., 2009). Given that action potential widths varied between ∼ 1.0 and 1.5 ms and assuming an absolute refractory period of ∼ 0.5-1.0 ms, the minimum ISI of a single neuron’s spike train should be ∼ 2 ms. A threshold of < 1 % of ISIs < 2 ms was therefore utilized for the inclusion of a putative single unit in this study (Mallet, Pogosyan, Marton, et al., 2008; Mallet, Pogosyan, Sharott, et al., 2008; Sharott et al., 2017). Of the 551 putative single unit spike trains reported here, the percentage of ISIs with durations < 2 ms was 0.0457, 0-0.227 %, suggesting minimal contamination by stray units.

All data were visually inspected in Neuroexplorer 4 (Nex Technologies, Colorado Springs, CO) and exported to MATLAB 2017b (MathWorks, Matick, MA) for analysis. Epochs with consistent, robust SWA or ACT were selected for analysis. During SWA, 30 second epochs were used to assess baseline neuronal activity. The impact of optogenetic inhibition during SWA was assessed by comparing neuronal activity 5 seconds prior to Arch-GFP stimulation with neuronal activity during 5 seconds of Arch-GFP stimulation. Five seconds of neuronal activity prior to (during SWA) and during tail pinch-evoked cortical ACT for 5 seconds were also compared. Neurons that demonstrated significant and consistent responses to Arch-GFP stimulation are reported. Neurons were deemed responsive if the firing rate or CV was consistently altered by at least 2 SD. PV-like GPe neurons were “isolated” from unidentified populations of GPe neurons in A2A-cre and GABRR3-cre animals by restricting analysis of unidentified GPe neurons to those with an in-:anti-phase ratio within the interquartile range of PV GPe neurons that were identified in PV-cre mice by optogenetic inhibition.

Mean firing rates were calculated from the number of spikes divided by epoch length. The CV was used as a metric of regularity. To examine the relationship between cortical SWA and neuronal firing, phase histograms were generated in MATLAB. SWA was extracted from the raw EEG signal by applying a bandpass 0.5-1.5 Hz 2^nd^ order Butterworth filter in the forward and reverse directions to avoid phase shifts. The EEG was downsampled to 1 kHz using the MATLAB function “resample”, then the instantaneous phase of the EEG was calculated from the Hilbert transform (Le Van Quyen et al., 2001). In order to correct for the non-sinusoidal nature of slow cortical oscillations, the empirical cumulative distribution function (MATLAB) was applied (Abdi et al., 2015; Mallet, Pogosyan, Sharott, et al., 2008; Siapas, Lubenov, & Wilson, 2005). Thus, each spike was assigned to a phase of the EEG from 0-360° (0°/360° and 180° corresponding to the peak active and inactive components of the EEG, respectively). Data were binned at 10° in figures, and at 180° for statistical comparisons. Each neuron’s spike probability was calculated from (spikes/bin)/(total # of spikes) X 100%. Population phase histograms are plotted as median and interquartile range. The multitaper Fourier transform function (Bokil, Andrews, Kulkarni, Mehta, & Mitra, 2010; Brazhnik, McCoy, Novikov, Hatch, & Walters, 2016; McConnell, So, Hilliard, Lopomo, & Grill, 2012) was applied using MATLAB to assess spectral power within the downsampled 1 kHz EEG signal (chronux.org; NW = 3, K = 5). 5s epochs during SWA or ACT were examined to determine total power in the 0.5-1.5 Hz, 10-39.9 Hz, and 40-250 Hz frequency bands respectively. Power within each band of interest was then normalized to the total power from 0-250Hz to control for variability in signal amplitude between recordings.

### Ex vivo electrophysiological recording

Acute brain slices were prepared from B6.129P2-Pvalb^tm1(cre)Arbr^/J X B6.Cg-Gt(ROSA)26Sor^tm9(CAG-tdTomato)Hze^/J mice as previously described (Chazalon et al., 2018). Briefly, animals were anesthetized with ketamine/xylazine (100/20 mg/kg, respectively) and perfused transcardially with ice-cold modified artificial cerebrospinal fluid (ACSF), equilibrated with 95% O_2_ and 5% CO_2_, and containing (in mM): 230 sucrose, 26 NaHCO_3_, 2.5 KCl, 1.25 NaH_2_PO_4_, 0.5 CaCl_2_, 10 MgSO_4_ and 10 glucose. Brains were rapidly removed and sectioned into 300 μm-thick parasagittal slices with a vibrating blade microtome (VT1200S; Leica Microsystems, Germany). Slices containing the GPe were then left to equilibrate for 1 h (at 35°C) in ACSF of the following composition (in mM): 126 NaCl, 26 NaHCO_3_, 2.5 KCl, 1.25 NaH_2_PO_4_, 2 CaCl_2_, 2 MgSO_4_, 10 glucose, 1 sodium pyruvate and 4.9 reduced L-gluthathione (equilibrated with 95% O_2_–5% CO_2_). Individual brain slices were placed in a recording chamber where they were perfused at 4–5 ml/min with synthetic interstitial fluid (SIF) at 35 °C containing (in mM): 126 NaCl, 3 KCl, 1.25 NaH_2_PO_4_, 1.6 CaCl_2_, 1.5 MgSO_4_, 10 glucose and 26 NaHCO_3_ (equilibrated with 95 % O_2_ and 5 % CO_2_). Somatic patch-clamp recordings were obtained under visual guidance (E600FN Eclipse workstation, Nikon, Japan; Nikon Fluor 60 X/1.0 NA) using motorized manipulators (Patchman NP2, Eppendorf, France). PV GPe neurons were identified by visualization of tdTomato under epifluorescence. Autonomous PV GPe neuron activity was recorded in the presence of the GABA_A_ receptor (GABAzine, 20 µM), GABA_B_ receptor (CGP 55845, 1 µM), AMPA/Kainate receptor (DNQX, 20 µM), and NMDA receptor (D-APV, 50 µM) antagonists in the loose-seal configuration in current clamp mode using borosilicate glass pipettes (4-6 MΩ) containing (in mM): 140 NaCl, 23 glucose, 15 HEPES, 3 KCl, 1.5 MgCl_2_, 1.6 CaCl_2_. pH and osmolarity were adjusted to 7.2 with 1 M NaOH and to 300-310 mOsm, respectively. Electrophysiological recordings were acquired using a computer running Clampex 9.2 software (Molecular Devices, Palo Alto, CA, USA) connected to a Multiclamp 700B amplifier (Molecular Devices) via a Digidata 1320A digitizer (Molecular Devices). Data were low-pass filtered at 4 kHz and sampled at 20 kHz.

### Histological processing of ex vivo tissue

B6.129P2-Pvalb^tm1(cre)Arbr^/J X B6.Cg-Gt(ROSA)26Sor^tm9(CAG-tdTomato)Hze^/J mice were euthanized with 20 % urethane and transcardially perfused with PBS, followed by 4 % w/v paraformaldehyde in 0.1 M PB, pH 7.4. The brain was then removed and incubated overnight in the same fixative at 4°C, then immersed in PBS containing 20 % w/v sucrose for 24h at 4°C, and stored in this solution at −80°C before being sectioned in the coronal plane at 50 μm on a cryostat (CM3000; Leica Microsystems Inc.). Sections were then washed in PBS and those containing ‘rostral’, ‘central’, and ‘caudal’ GPe (corresponding approximately to AP −0.2 mm, −0.45 mm, and −0.7 mm from Bregma, respectively (Paxinos & Franklin, 2001) were selected for immunohistochemical detection of GPe markers. Sections were incubated overnight at RT in a mixture of 3 primary antibodies diluted in PBS-T (1:100 goat anti-FoxP2 P2; 1:100; sc-21069; Santa Cruz; 1:1,000 guinea pig anti-parvalbumin; 195004; Synaptic Systems; 1:1,000 rat anti red fluorescent protein 5f8; Chromotek). Sections were then washed and incubated for 1 hour at RT in PBS-T containing a mixture of secondary antibodies (1:500 Alexa Fluor 488 donkey anti guinea-pig IgG; A11073; Life Technologies; 1:500 DyLight 594 donkey anti rat IgG; NBP1-75661; Novus; 1:500 Alexa Fluor 647 donkey anti-goat IgG; A21447; Life Technologies). Finally, sections were washed in PBS, mounted in Vectashield (Vector Laboratories), and imaged on a confocal fluorescence microscope (TCS SP8, Leica Microsystems Inc.). Images were acquired using a 20 X 1.8 NA objective lens in 1.0 µm steps between 2 µm and 17 µm from the upper surface of each section. Colocalization was assessed from maximal z-projection images using the cell counter plug-in of ImageJ.

Sections containing the striatum were processed for the immunohistochemical detection of TH. Sections were first incubated in primary antibody (1:10000 monoclonal anti-TH; MAB318, Millipore Bioscience Research Reagents) in PBS-T overnight at RT. Subsequently, the sections were incubated in secondary antibody in PBS-T (1:1000 biotinylated horse anti-mouse IgG; Vector Laboratories) for 90 min at RT. Finally, sections were incubated in avidin-biotin peroxidase complex (1:500; Vector Laboratories) for 60 min at RT and immunoreactivity was revealed using AMEC (Vector Laboratories). Sections were then rinsed, mounted on gelatin-coated slides, and coverslipped in VectaMount (Vector Laboratories). At least three sections from each hemisphere containing the striatum were scanned in an Epson expression 10000XL high-resolution scanner. Mean optical density was measured in the top half of the striatum (Mercator; Explora Nova, La Rochelle, France) and values were corrected for background staining as above. TH immunoreactivity ipsilateral to vehicle or 6-OHDA injection was expressed as a percentage of immunoreactivity in the contralateral hemisphere, as described above (vehicle: ipsilateral striatal TH immunoreactivity = 99, 99-131 %; n = 3; 6-OHDA: ipsilateral striatal TH immunoreactivity = −15, −29 −6 %; n = 7; p = 0.01667; MWU).

## Acknowledgements

This study was funded by NIH NINDS grants 2R37 NS041280, P50 NS047085, and 5T32 NS041234. Confocal imaging work performed at the Northwestern Center for Advanced Microscopy, supported by NCI CCSG grant P30 CA060553.

---

**Figure S1.**
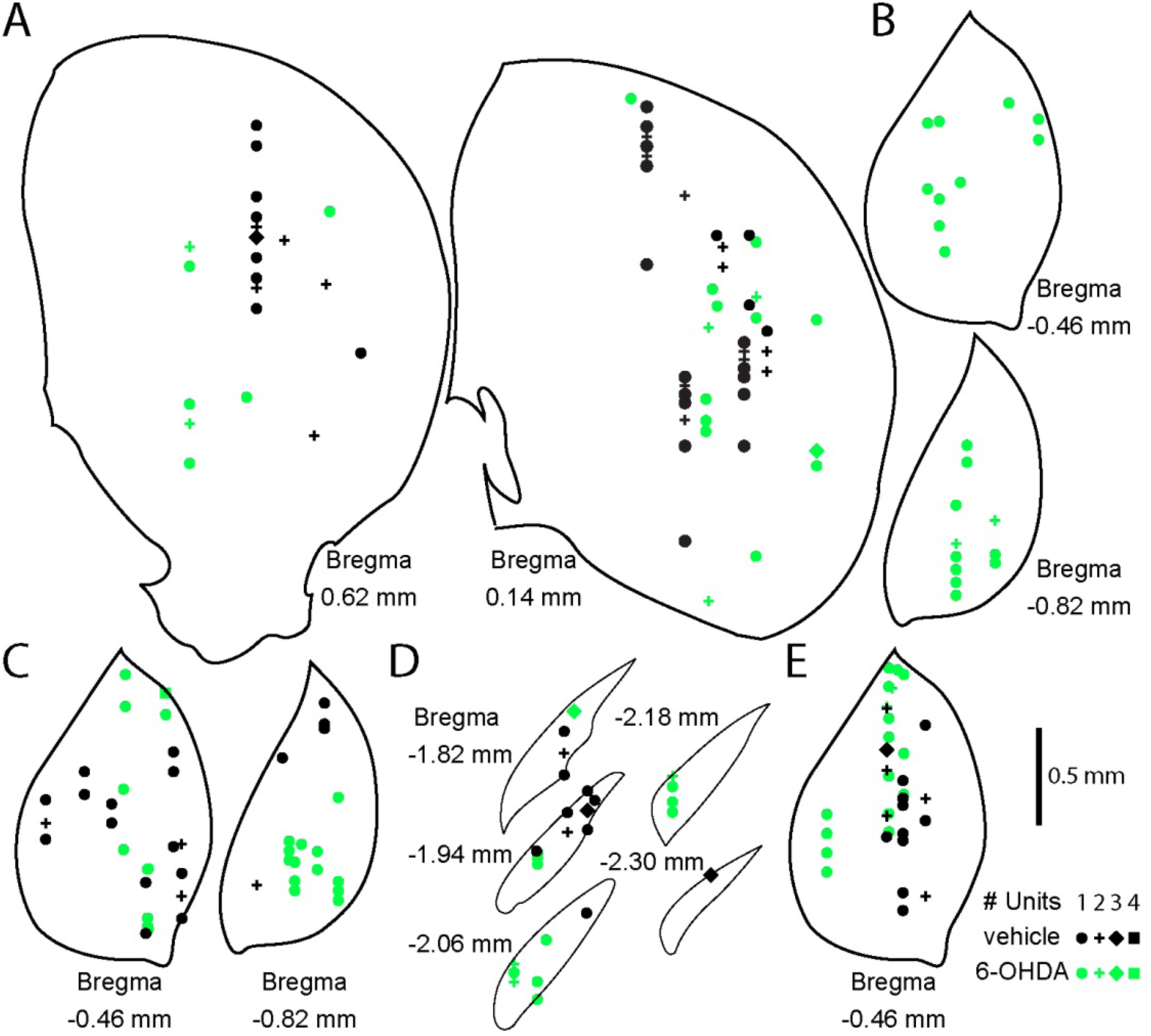
Recording site map of recorded units. (A-E) Map of electrode recording sites used in this study. (A) Striatal electrode sites from A2A-cre mouse recordings. (B) GPe electrode sites from A2A-Cre mouse recordings. (C) GPe electrode sites from PV-Cre mouse recordings. (D) STN recording sites from PV-Cre mouse recordings. (E) GPe electrode sites from GABRR3-Cre mouse recordings. Vehicle- and 6-OHDA-injected mouse recording sites plotted as black and green, respectively. Number of units per site indicated by marker shape (see key in, E).

**Table S1.** Spectral analysis of EEG during SWA and ACT.

| Frequency Band (Hz) | Normalized Total Power in Control | Normalized Total Power in 6-OHDA | p - value | Peak Frequency in Control (Hz) | Peak Frequency in 6-OHDA (Hz) | p - value | n's (Control/ 6-OHDA) |
| --- | --- | --- | --- | --- | --- | --- | --- |
| 0.5-1.5 (SWA) | 0.247, 0.169-0.375 | 0.233, 0.124-0.335, | 0.2769 | 1.22, 0.855-1.34 | 1.1, 0.855-1.40 | 0.7042 | 60/69 |
| 10-39.9 (SWA) | (0.7919, 0.53-1.23) $\times 10^{-2}$ | (0.827, 0.617-1.21) | 0.6455 | 10.4, 10.1-11.4 | 10.9, 10.3-12.3 | 0.0121* | 60/69 |
| 40-250 (SWA) | (0.332, 0.235-0.565) $\times 10^{-2}$ | (0.341, 0.193-0.523) $\times 10^{-2}$ | 0.4576 | 48.3, 44.1-54.1 | 45.0, 41.9-49.7 | 0.0090* | 60/69 |
| 0.5-1.5 (ACT) | (6.97, 3.43-13.4) $\times 10^{-2}$ | (7.21, 4.39-13.4) $\times 10^{-2}$ | 0.9121 | 0.610, 0.610-1.22 | 0.610, 0.610-1.47 | 0.7522 | 39/35 |
| 10-39.9 (ACT) | (2.55, 0.798-4.15) $\times 10^{-2}$ | (2.51, 1.88-4.18) $\times 10^{-2}$ | 0.2175 | 11, 10.1-12.7 | 11.5, 10.5-14.7 | 0.2807 | 39/35 |
| 40-250 (ACT) | (1.39, 0.608-2.44) $\times 10^{-2}$ | (1.47, 0.674-2.11) $\times 10^{-2}$ | 0.9549 | 45.7, 41.8-51.5 | 42.7, 41.0-44.7 | 0.0103* | 39/35 |

Total power and peak frequency during SWA or ACT was assessed in the 0.5-1.5 Hz, 10-39.9 Hz, and 40-250 Hz frequency bands. Spectral power was normalized to total power in the 0-250 Hz range to account for variability in signal amplitude between recordings. \*, $p < 0.05$ (MWU).

**Table S2.**
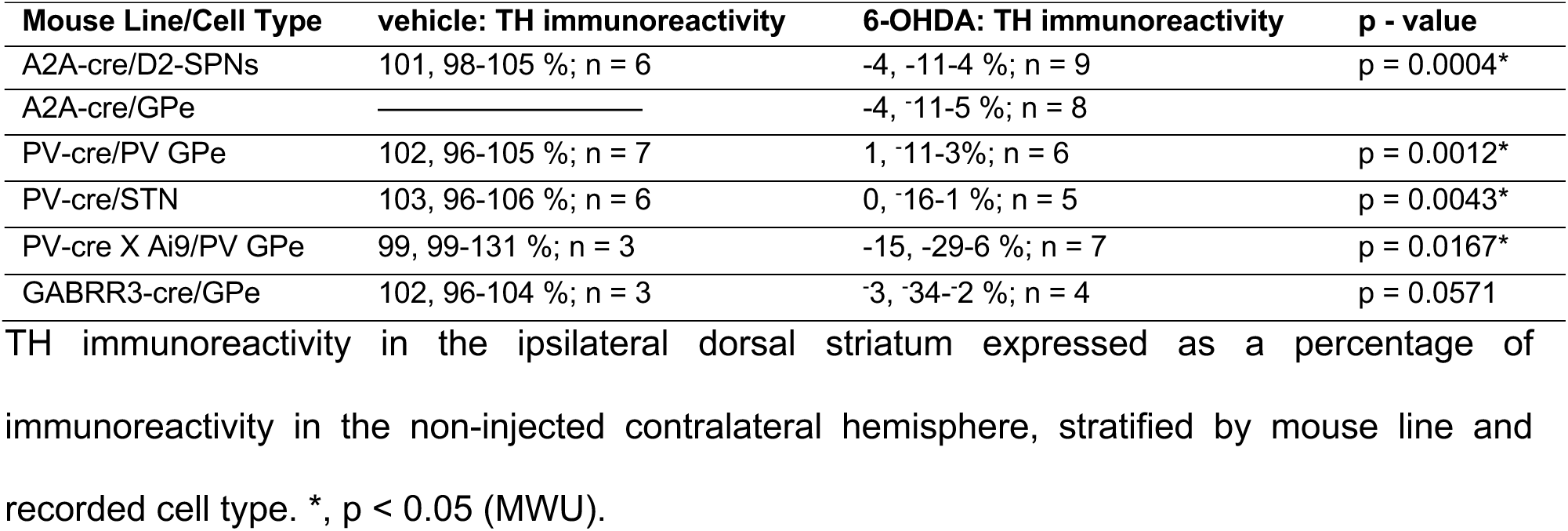
Tyrosine Hydroxylase Immunoreactivity.

